# Multi-resolution imaging using bioluminescence resonance energy transfer identifies distinct biodistribution profiles of extracellular vesicles and exomeres with redirected tropism

**DOI:** 10.1101/2020.03.27.012625

**Authors:** Anthony Yan-Tang Wu, Yun-Chieh Sung, Yen-Ju Chen, Steven Ting-Yu Chou, Vanessa Guo, Jasper Che-Yung Chien, John Jun-Sheng Ko, Alan Ling Yang, Hsi-Chien Huang, Ju-Chen Chuang, Syuan Wu, Meng-Ru Ho, Maria Ericsson, Wan-Wan Lin, Chantal Hoi Yin Cheung, Hsueh-Fen Juan, Koji Ueda, Yunching Chen, Charles Pin-Kuang Lai

## Abstract

Extracellular particles (EP) including extracellular vesicles (EVs) and exomeres have been shown to play significant roles in diseases and therapeutic applications. However, their spatiotemporal dynamics *in vivo* have remained largely unresolved in detail due to the lack of a suitable method. We therefore created a bioluminescence resonance energy transfer (BRET)-based reporter, PalmGRET, to enable pan-EP labelling ranging from exomeres (< 50 nm) to small (< 200 nm) and medium and large (> 200 nm) EVs. PalmGRET emits robust, sustained signals and allows the visualization, tracking and quantification of the EPs from whole-animal to nanoscopic resolutions under different imaging modalities, including bioluminescence, BRET and fluorescence. Using PalmGRET, we show that EPs released by lung metastatic hepatocellular carcinoma (HCC) exhibit lung tropism with varying distributions to other major organs in immunocompetent mice. We further demonstrate that gene knockdown of lung-tropic membrane proteins, solute carrier organic anion transporter family member 2A1 (Slco2a1), alanine aminopeptidase (Cd13) and chloride intracellular channel (Clic1) decreases HCC-EP distribution to the lungs and yields distinct biodistribution profiles. We anticipate that EP-specific imaging, quantitative assays and detailed *in vivo* characterization to be a starting point for more accurate and comprehensive *in vivo* models of EP biology and therapeutic design.

## Introduction

Extracellular vesicles (EVs) and exomeres are extracellular particles (EP; 30–10,000 nm) released by cells to deliver bioactive cargoes to enable communication over short and long distances, both within and between organisms^[1, 2]^. EV subtypes include exosomes (30–100 nm in diameter), shed microvesicles (or ectosomes, microparticles; 100–1,000 nm) and apoptotic bodies (50–4,000 nm)^[3]^. The subtypes are lipid-bilayer encapsulated and released through two main pathways: i) fusion of the multivesicular bodies with the plasma membrane to release exosomes; or ii) outward budding and fission of the plasma membrane to release microvesicles. As EV subtype-specific isolation methods and terminology remain to be standardized^[4]^, the generic term “EV” is used here unless otherwise specified. Using asymmetric flow field-flow fractionation (AF4), cell-released EPs can be further resolved into Exo-S (60–80 nm), Exo-L (90–120 nm) small EVs (sEVs) and exomeres (< 50 nm), with the latter considered as non-EVs as they are non-membranous and exhibit a limited set of membrane-bound proteins^[4, 5]^.

To facilitate intercellular communication, EVs and exomeres ferry bioactive cargoes between cells. These include DNA, RNA (*e.g.* mRNA, miRNA and lncRNA) and proteins (*e.g.* ligands and receptors), which have been associated with development, immune response, neurodegenerative diseases, developmental disorders and cancer progression^[6–12]^. Tumor exosomes have been shown to be organotropic in facilitating subsequent metastasis to the tropic organs^[13]^. Specific integrins have been found to redirect tumor exosomes to the lungs (ITGβ_4_) and liver (ITGβ_5_). Concurrently, EVs are being actively explored as a diagnostic medium for liquid biopsy, in addition to serving as an endogenous delivery vehicle for therapeutic applications. Specifically, EVs have been designed with cell-type- and cancer-targeting moieties to enable the targeted delivery of therapeutic EVs^[14]^. However, the spatiotemporal dynamics of EVs *in vivo* remains elusive. For example, while organotropic and cell-targeting EVs may improve delivery to a particular site, their distribution to and relationship with other non-targeted tissues at organ systems level is largely unresolved. Moreover, accurate quantification of EVs has been confounded by a lack of suitable dyes and technical limitations^[15]^. The biological understanding and therapeutic development of EVs would be greatly aided by the detailed characterization of EVs’ spatiotemporal properties *in vivo*.

Conventional EV-labelling methods mainly rely on fluorescent (FL) dyes. Organic dyes such as PKH dyes, 1,1′-dioctadecyl-3,3,3′3′-tetramethylindocarbocyanine perchlorate (DiI) and its derivatives or carboxyfluorescein succinimidyl ester (CFSE) have been widely used in the study of EVs due to their stable and high FL signals^[16, 17]^. However, lipophilic FL dyes were later identified to spontaneously form nanometer sized micelles (*i.e.* false-positives) and carry an extended *in vivo* half-life (*e.g.* PKH2: 5–8 days, PKH26: > 100 days), resulting in inaccurate spatiotemporal detection of EVs^[18, 19]^.

Bioluminescent (BLI) reporters have been used to label EVs for the *in vivo* imaging and pharmacokinetic analysis of administered EVs^[20, 21]^. However, the resolution is relatively poor when compared with that of FL dyes, and so conventional methods typically use BLI and FL reporters for *in vivo* and *in vitro* imaging, respectively. Bioluminescence resonance energy transfer (BRET) was developed to overcome some of these limitations^[22]^. Similar to the principle of fluorescence resonance energy transfer, BRET reporters consist of a pair of BLI and FL proteins conjugated in close proximity (30–70 Å) by a linker for intramolecular energy transfer. When treated with the substrate, BRET will emit both BLI and FL signals for detection. However, the use of BRET has been limited by BLI protein size, luminescence stability and intensity^[23]^. Nanoluc (Nluc) has recently overcome these obstacles to BRET; Nluc is a small (19 kDa), ATP-independent luciferase that provides one of the brightest glow-type bioluminescence for sustained excitation of the paired FL protein^[23–26]^.

Previous studies have used EV markers fused to FL proteins (*e.g.* CD63-GFP) to examine EV distributions, but have mostly been limited to *in vitro* observations^[27]^. Meanwhile, multiphoton microscopy enables high-resolution imaging of cells *in vivo* by exciting fluorophores with multiple photons at longer wavelengths for deeper tissue penetration (up to ~1 mm)^[28]^. However, it generally requires the imaging window to be fixed at one location to record cellular events longitudinally. We applied multiphoton microscopy to achieve *in vivo* visualization of EV release using pan-EV labelling FL EV reporters^[29]^. However, as EVs are circulated, they cannot be systemically tracked by multiphoton microscopy. Multiharmonic microscopy has also been applied successfully to imaging EVs *in vivo*, but it relies on the presence of specific metabolite signatures, and so is inapplicable to unspecified EVs^[30, 31]^. On the other hand, radioactive tracers such as ^99m^Tc-HMPAO and ^111^In-oxine have been applied to label EVs for *in vivo* imaging and EV biodistribution analysis using positron emission tomography–computed tomography and single-photon emission computed tomography, respectively^[32, 33]^. However, the use of radioisotopes is limited to well-trained operators and certified facilities with available instruments, making these techniques costly and unsuitable for a general laboratory.

To overcome these limitations of EV visualization, we describe here a BRET-based EP reporter, PalmGRET, that enables multimodal, multi-resolution imaging and quantification of EPs from organismal to super-resolutions. PalmGRET was created by fusing a palmitoylation signal peptide sequence from growth-associated protein 43 (GAP43)^[34]^ to the N-terminus of a GFP-Nluc BRET reporter (GpNluc)^[23]^. We demonstrate that PalmGRET can label multiple EP populations, including exomeres (< 50 nm) and small (< 200 nm; sEVs), medium and large (> 200 nm; mEVs and lEVs) EVs, making it a versatile reporter to visualize and monitor EPs released by cells. Moreover, PalmGRET specifically labels the EV inner membrane with a long-term and robust signal, thereby enabling live cell BLI and BRET microscopy, as well as super-resolution radial fluctuations (SRRF) nanoscopy^[35]^ to observe EP dynamics *in vitro*. With its emitted BLI and BRET-FL signals, PalmGRET further allows *in vivo* imaging of administered EPs over an extended period of time (~20 min), as well as highly sensitive EP biodistribution analysis to detect minute EP changes in organs. Using PalmGRET, we identified lung-tropic proteins of EPs derived from lung metastatic hepatocellular carcinoma (HCC) in immunocompetent C3H mice, Slco2a1 (solute carrier organic anion transporter family member 2A1, Cd13 (Anpep, alanine aminopeptidase), and Clic1. Knockdown of the lung-tropic proteins significantly reduced HCC EP lung tropism. Additionally, Slco2a1 and Cd13 protein knockdowns of HCAC EPs resulted in redirection to the kidney and heart. Our results suggest that the *in vivo* function and therapeutic design of EPs should be considered at organ system level to account for their dynamic biodistribution.

## Results and Discussion

### PalmGRET labels EVs at the inner membrane with FL and BLI reporter activities

To develop a multimodal and multi-resolution EP reporter, PalmGRET was created by genetically fusing the palmitoylation signal peptide sequence^[36]^ of GAP43^[34]^ to the N-terminus of a BRET pair, GpNluc^[23]^ (**Figure 1 a–c**). Upon stable expression in human 293T cells, PalmGRET uniformly labelled the plasma membrane to reveal an uneven membrane surface with bud-like structures at the apical plane and protrusions of the cells, which may be precursors to EP release (**Figure 1d**). To examine whether PalmGRET can be applied to visualize EP release from multiple cellular populations, stable 293T-PalmGRET and 293T-PalmtdTomato cells^[29]^ were co-cultured; both cell types were found to demonstrate EP release, thereby suggesting bidirectional EP-mediated intercellular communication (**Figure 1e and Movie S1**).

**Figure 1.**
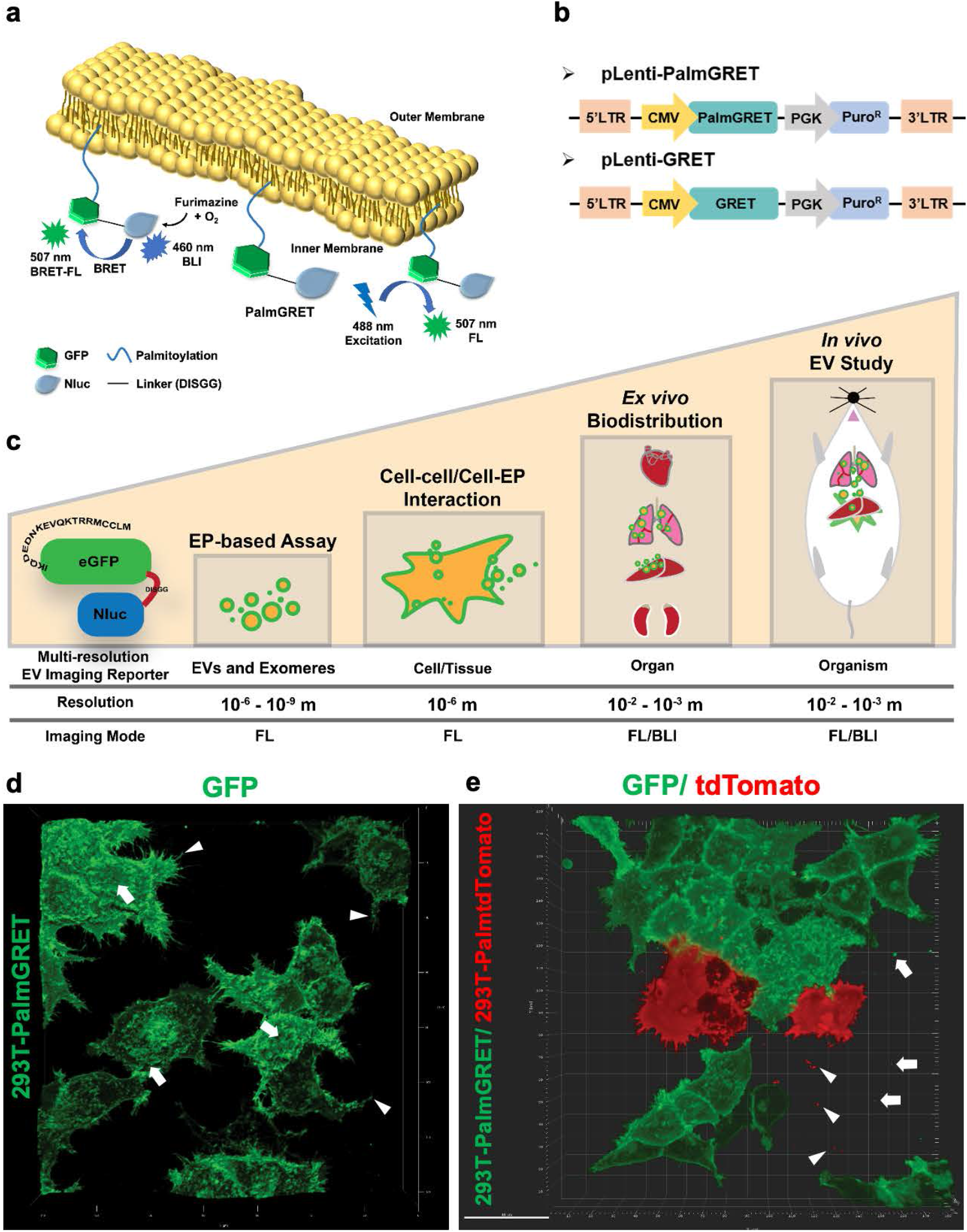
PalmGRET labels cellular and EV membranes. **(a)** Representation of multimodal PalmGRET labelling the inner leaflet of the plasma membrane of cells and EVs to emit BLI and FL using BRET excitation (BRET-FL), as well as FL from external light excitation. **(b)** Lentiviral constructs for PalmGRET and GRET (control) for EP labelling. **(c)** Multi-resolution EP imaging. PalmGRET labels cellular membranes to enable imaging of EV donor cells and EVs from nanoscopic to macroscopic resolutions by FL and/or BLI signals. **(d)** 3D reconstruction of Z-stack, live-cell confocal micrographs of 293T-PalmGRET cells, revealing uneven, budding membrane surfaces (arrows) and protrusions (arrowheads) that may be precursors of EV release. **(e)** Co-culture of 293T-PalmGRET and 293T-PalmtdTomato cells showing released EPs in the extracellular space. 293T-PalmGRET EPs (arrows); 293T-PalmtdTomato EPs (arrowheads). Bar, 40 μm.

To characterize the sEV labelling properties of PalmGRET, sEVs isolated from stable 293T-PalmGRET and 293T-GRET cells (control) were examined by nanoparticle tracking analysis. The resulting sEVs, sEV-PalmGRET (147.7 ± 2.2 nm), sEV-GRET (143.9 ± 3.8 nm; control) and sEV-wildtype (EV-WT; 150.0 ± 3.5 nm; control), were all similar in size (**Figure 2a, b**). Transmission electron microscopy with immunogold labelling further demonstrated that PalmGRET mostly localizes to sEV membranes (**Figure 2c, d**). To mitigate possible steric hindrance to ligand–receptor interaction on the outer EV membrane ^[13, 37]^, we designed PalmGRET to label the inner EV membrane using the palmitoylation signal. To confirm the inner-membrane specificity of PalmGRET, we subjected the labelled sEVs to dot blot analysis in the presence or absence of detergent (**Figure 2e, f**). Whereas antibodies can only recognize proteins on the outer sEV membrane in the absence of the detergent, the addition of TWEEN-20 (0.1% v/v) enables antibody entry and immunolabeling of proteins at the inner membrane by disrupting the EV membranes. Therefore, as sEV-GRET showed a dim signal with the detergent and no signal in its absence, GRET appeared to be non-specifically packaged into the EV lumen (**Figure 2e**). By contrast, a strong sEV-PalmGRET signal was detected only in the presence of the detergent, thereby indicating inner membrane-specific labelling of sEVs (**Figure 2f**).

**Figure 2.**
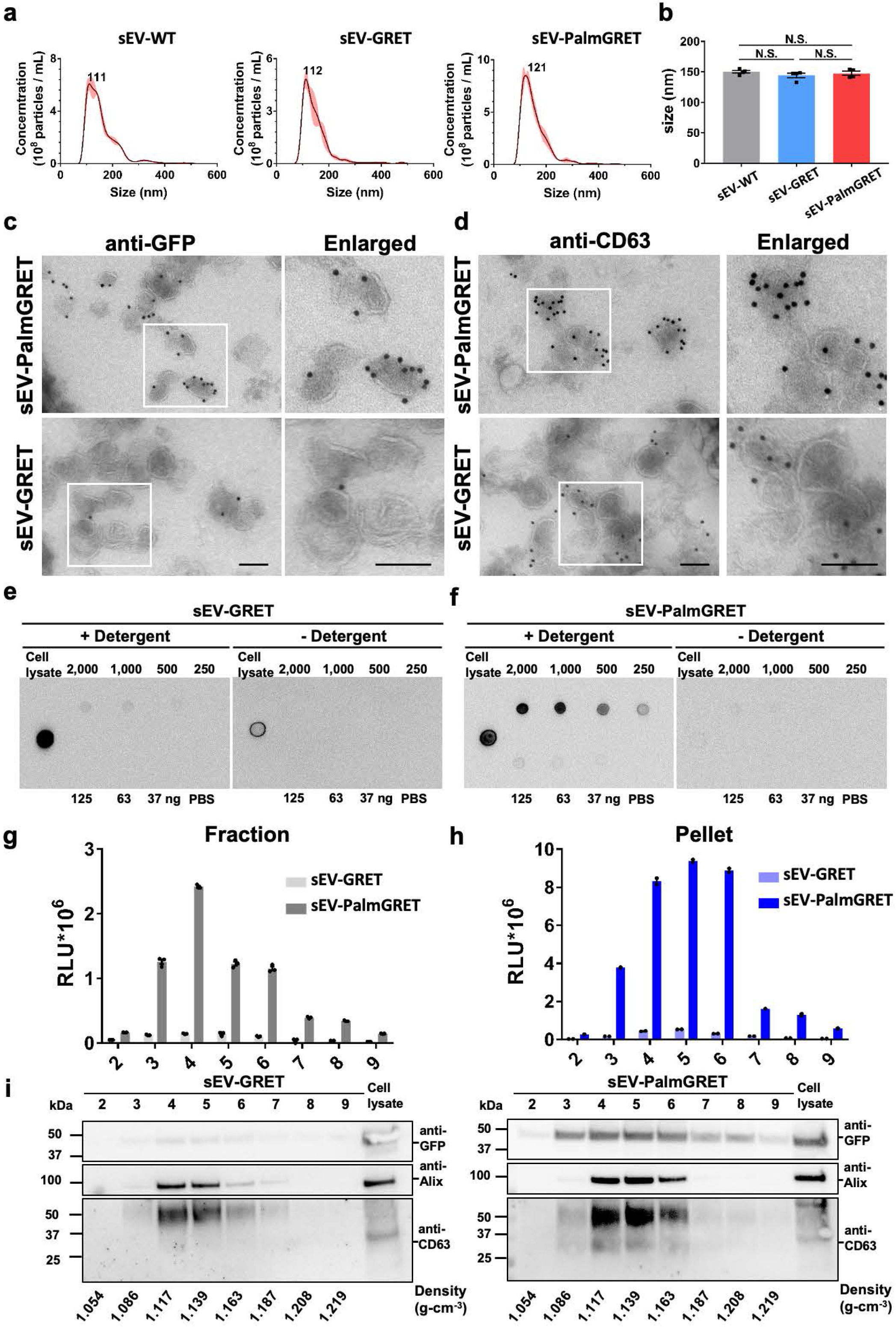
PalmGRET labels inner EV membrane and exhibits robust BLI signals in EVs. **(a, b)** Nanoparticle tracking analysis of sEV-wildtype (sEV-WT), sEV-GRET and sEV-PalmGRET, demonstrating their similar **(a)** size distributions (peaks at 111, 112 and 121 nm, respectively; representative data of three independent experiments) and **(b)** mean sizes (determined from four independent experiments). One-way ANOVA followed by Tukey’s *post hoc* test was used. N.S., non-significant with *p* > 0.05. **(c, d)** Transmission electron micrographs showing that PalmGRET labels sEVs at their membranes. Micrographs of isolated sEV-293T-PalmGRET and EV-293T-GRET immunogold labelled with **(c)** anti-GFP or **(d)** anti-CD63, and secondary gold-conjugated secondary antibodies. Bar, 100 nm. **(e, f)** Dot blot analysis showing that PalmGRET labels the sEV inner membrane. A strong PalmGRET signal was only detected in the presence of **(f)** Tween-20 detergent [0.1% (v/v)] in a dose-dependent manner when compared with **(e)** the GRET control, thereby indicating inner EV membrane-specific labelling by PalmGRET. Cell lysates were applied as positive controls. **(g, h)** Nluc activity assay of sEV-293T-PalmGRET and sEV-293T-GRET following **(g)** sucrose gradient fractionation and **(h)** fraction pelleting. A significantly higher Nluc activity was detected in sEV-containing fractions (Nos. 3–6) of sEV-PalmGRET-EV when compared with sEV-293T-GRET. **(i)** Western blot analysis of sEV proteins isolated from the pelleted fractions revealed that PalmGRET (right), but not GRET (left), coincides with Alix (95 kDa) and CD63 (30–60 kDa). Cell lysates of 293T-PalmGRET and 293T-GRET were applied as positive controls. The expected sizes of PalmGRET and GRET are 49 and 46 kDa, respectively.

To verify the BLI function and EV-specificity of PalmGRET, isolated sEVs were subjected to sucrose density gradient fractionation followed by BLI assay. In both the sEV fractions (**Figure 2g**) and pelleted sEVs (**Figure 2h**), sEV-PalmGRET showed ~20 times the BLI activity of the sEV-GRET control. Western blot analysis of the pelleted sEVs further demonstrated that PalmGRET coincided with sEV markers (CD63 and Alix)^[38]^ at the sEV-containing fractions (1.117–1.163 g·cm^−3^; **Figure 2i** and **Figure S1**). Overall, PalmGRET efficiently and specifically labelled the inner sEV membrane while functioning as a BLI reporter.

### PalmGRET labels multiple bionanoparticle populations

An EV subtype-specific protein remains to be identified, but multiple tetraspanins (CD63, CD81 and CD9) have been recommended to identify subpopulations of sEVs^[39]^. While studies have commonly focused on a specific EV subpopulation to investigate its biological function, we envision multiple EP populations working simultaneously in mediating EP-based intercellular communication, especially *in vivo*. Therefore, we designed PalmGRET as a pan-EP labelling reporter to label and examine multiple EP populations. Using AF4, Zhang *et al.* subcategorized sEVs as Exo-L (90–120 nm), Exo-S (60–80 nm), and identified non-membranous exomeres (< 50 nm), each with different molecular profiles^[40]^. To further examine the pan-labelling coverage of EP by PalmGRET, we performed AF4 on sEV-PalmBRET and characterized fractions corresponding to Exo-L (P4), Exo-S (P3) and exomeres (P2) (**Figure 3a**). To verify PalmGRET-labelling of the sEV and exomere fractions, the fractions (P2, P3 and P4) were subjected to BLI assay, which revealed ~10-fold stronger BLI and BRET-FL signals in PalmGRET-labelled Exo-L, Exo-S and exomeres when compared to that of GRET (**Figure 3b, c**). While Exo-S and Exo-L of EV-PalmGRET exhibited no significant differences, exomeres showed significantly lower signals in both BLI and BRET-FL activities, which positively correlates to the particle abundance of the EPs as found by the quasi-elastic light scattering detector (**Figure 3a**). Interestingly, the PalmGRET-labelled exomere fraction showed a slightly greater hydrodynamic radius (~40 nm) than the GRET control (~35 nm), suggesting that PalmGRET slightly increases the size of exomeres while remaining under 50 nm in diameter. We further demonstrated that PalmGRET effectively labels EVs isolated from 0.22, 0.8 and 1.2 μm-filtered media, as well as 10k and 100k pellets (**Figure S2, S3**). We therefore confirmed that PalmGRET labels varying sized EPs ranging from exomeres, and sEVs to lEVs.

**Figure 3.**
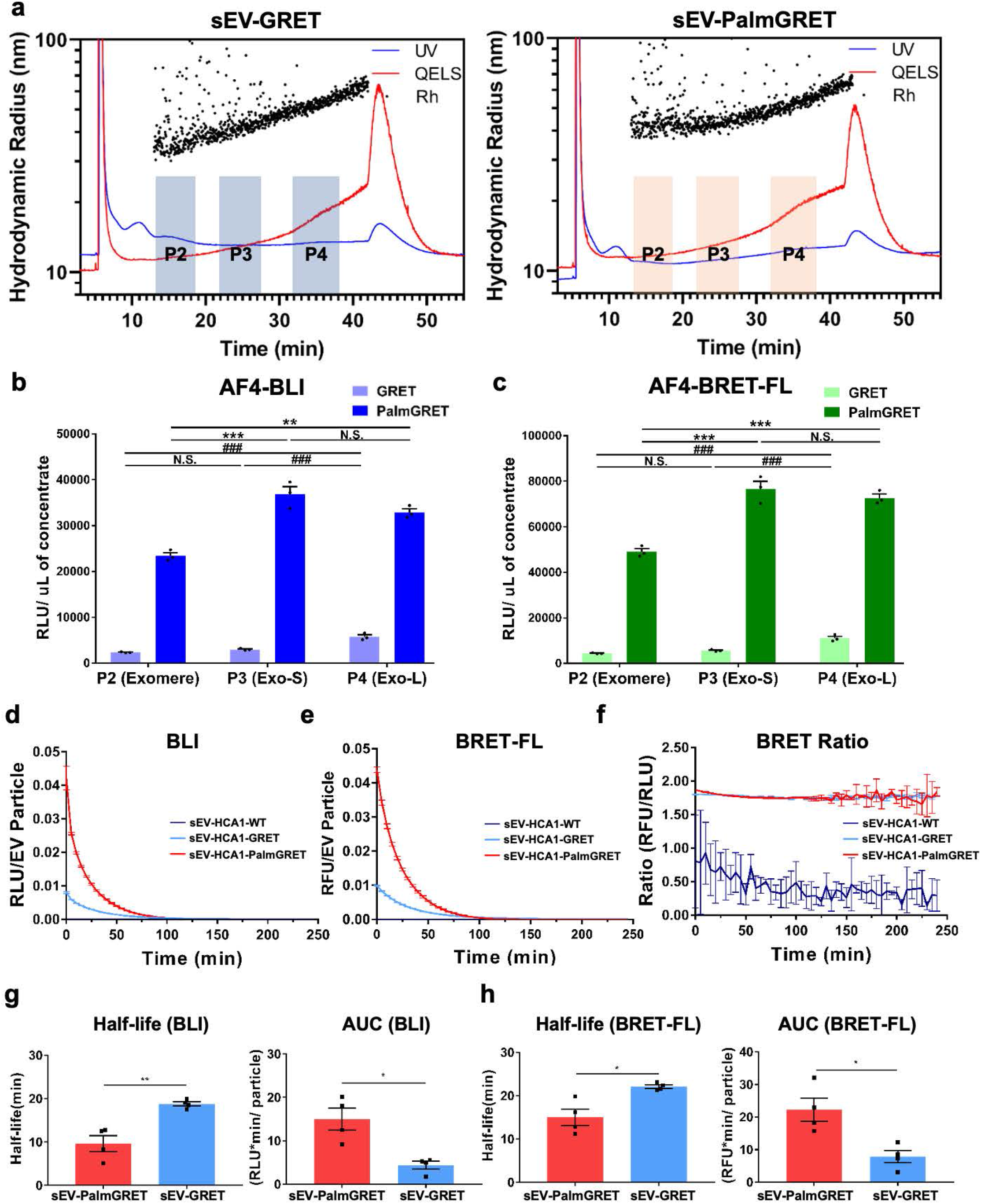
PalmGRET labels multiple EP populations and exhibits long-term reporter signals in EVs. **(a)** AF4 analysis of sEVs isolated from stable 293T-PalmGRET and 293T-GRET cells. PalmGRET appears to increase the size of exomeres (P2) but not Exo-S (P3) or Exo-L (P4) when compared with the control (GRET). Scattered dots represent the hydrodynamic radii of detected particles. The blue solid line indicates relative particle abundance detected using UV (280 nm) light. The red solid line represents particle abundance detected using quasi-elastic light scattering. **(b, c)** Nluc and BRET activities of AF4 fractionated bionanoparticles indicating pan-labelling by PalmGRET. Thus labelled bionanoparticles showed 10-times higher **(b)** BLI and **(c)** BRET-FL signals in exomere, Exo-S and Exo-L fractions than the control (GRET). The resulting BLI and BRET-FL activities of different subpopulations were normalized to the volume of concentrate from the Amicon Ultra-15 30-kDa molecular weight cut-off concentration. N.S., *p* > 0.05; **/##, *p* < 0.01; ***/###, *p* < 0.001 with one-way ANOVA followed by Tukey’s *post hoc* test. **(d–h)** Nluc and BRET activity analyses revealed that sEV-PalmGRET emits strong and sustained **(d)** BLI and **(e)** BRET-GFP signals. **(f)** Both sEV-293T-PalmGRET and sEV-293T-GRET exhibited a similar BRET ratio, suggesting that BRET efficiency is not affected by the palmitoylation moiety in sEV-293T-PalmGRET. While sEV-GRET showed a longer half-life, sEV-PalmGRET exhibited greater AUCs in its emitted **(g)** Nluc and **(h)** BRET-FL signals during four independent experiments. **RFU**, relative fluorescence unit; **RLU**, relative light unit. *, *p* < 0.05; **, *p* < 0.01 with two-tailed Student’s t-test.

### PalmGRET-labelled EPs exhibit robust and long-term BLI and BRET-FL signals

To characterize PalmGRET’s BLI properties, sEV-PalmGRET was supplemented with furimazine (Fz) substrate and measured for BLI and BRET-FL activities over time. sEV-PalmGRET demonstrated a significantly higher signal in both the BLI and BRET-FL channels than either sEV-GRET or sEV-WT control (**Figure 3d, e**). Interestingly, 293T-GRET cells emitted higher BLI and BRET-FL signals than 293T-PalmGRET cells (**Figure S4a, b**). sEV-PalmGRET showed a similar BRET ratio to sEV-GRET, indicating that the labelling of GRET to sEVs using the palmitoylation moiety does not affect BRET efficiency (**Figure 3f**). The BRET ratio of 293T-PalmGRET and 293T-GRET cells were also similar, further supporting that PalmGRET does not affect reporter function (**Figure S4c**). While sEV-GRET exhibited approximately twice the BLI signal half-life of sEV-PalmGRET (18.8 ± 0.99 *vs*. 9.7 ± 3.67 min, respectively) and a significantly longer BRET-FL signal half-life (22.1 ± 0.41 *vs*. 15.0 ± 1.93 min, respectively), sEV-PalmGRET displayed a distinctly higher signal over time (AUC, area under curve) than sEV-GRET to enable robust signals and prolonged imaging of EPs (BLI: 15.0 ± 2.51 *vs*. 4.4 ± 0.90 RLU·min/particles, respectively; BRET-BLI: 22.0 ± 3.60 *vs*. 7.9 ± 1.89 RFU·min/particles, respectively) (**Figure 3g, h**). In contrast, 293T-PalmGRET cells showed a longer half-life than 293T-GRET cells (BLI: 27.4 ± 1.30 *vs*. 21.3 ± 0.35 min, respectively; BRET-FL: 26.8 ± 1.4 *vs*. 19.7 ± 0.32 min, respectively) but lower AUCs (BLI: 20.7 ± 1.25 *vs*. 31.2 ± 0.66 RLU·min/particles, respectively; BRET-FL: 20 ± 1.30 *vs*. 28.2 ± 0.87 RLU·min/particles, respectively) (**Figure S4d, e**). The observed difference in PalmGRET activities between the labelled EVs and cells may be attributed to a variation in membrane-to-cytosol ratio: an EV particle has a higher membrane-to-cytosol ratio, which is higher for an EV particle than for a cell^[41]^. In this way, PalmGRET locates to the plasma membrane and hence is more enriched per EV, whereas GRET predominately distributes to the cytosol and therefore is more abundant per cell. Consequently, sEV-PalmGRET showed stronger BLI and BRET-FL signals (and lower half-lives) as more Nluc enzyme is available to oxidize Fz when compared with sEV-GRET, thereby yielding greater AUCs. Similarly, 293T-GRET cells generate higher AUCs with more abundant Nluc available per cell. Taken together, PalmGRET emits strong and sustained signals for extended visualization and tracking of EPs.

### PalmGRET enables multimodal and super-resolution imaging of cells and EPs

To examine PalmGRET imaging functions under microscopy, 293T-PalmGRET and 293T-GRET cells were examined for BRET-FL and FL signals under live-cell microscopy. We first confirmed BRET-FL activity by detecting the emitted GFP signal *via* BRET (BRET-FL) by bioluminescence microscopy following Fz substrate administration, where PalmGRET primarily locates to the membrane while GRET mostly accumulates at the cytosol (**Figure S5**). Epifluorescence microscopy was next applied to image the same samples, and similar subcellular distribution patterns of PalmGRET and GRET were observed, thereby validating the observed BRET-FL signals (**Figure 4a** and **Figure S5**). In addition, EPs could be readily visualized under epifluorescence microscopy only in 293T-PalmGRET and not 293T-GRET cells. To resolve EP signals at super-resolution, SRRF nanoscopy successfully resolved individual EP-PalmGRET signals (**Figure 4a,** i–ii). We therefore employed SRRF nanoscopy in all subsequent subcellular EV imaging experiments, and readily visualized released EPs at super-resolution in real-time *in vitro* (**Figure 4b, c** and **Movies S2 and S3**). We further tested the capability of PalmGRET in detecting EPs in fixed tissues, and successfully detected EP-PalmGRET signals inside cells of immunostained sections of EP-PalmGRET-injected mouse lung tissue (**Figure 4d**). These results demonstrate PalmGRET’s versatility in investigating cellular and subcellular distribution of EPs in live cells, as well as in preserved tissues.

**Figure 4.**
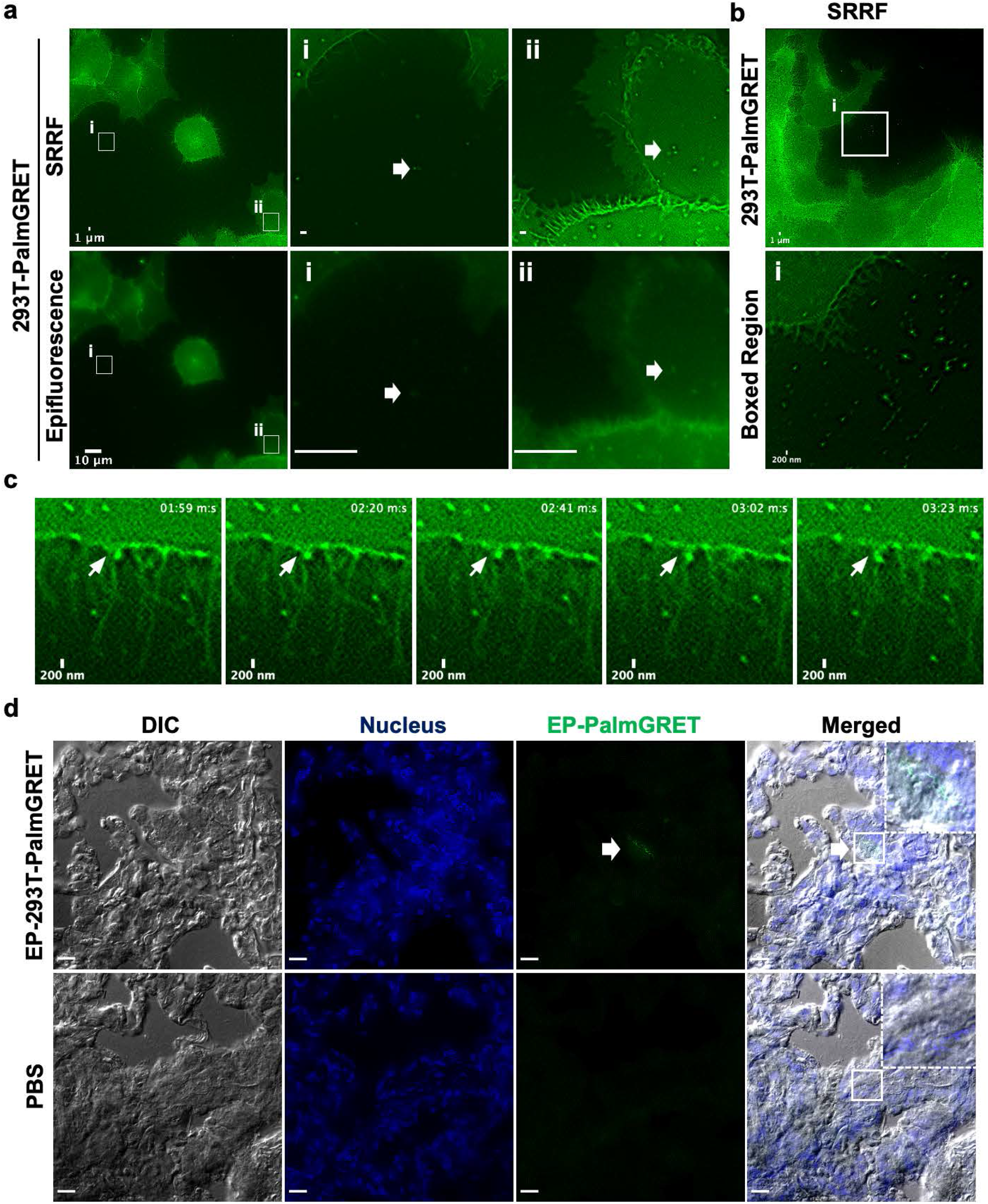
PalmGRET enables super-resolution fluorescence imaging of EPs *in vivo* and *ex vivo*. **(a)** Live-cell, fluorescence imaging of 293T-PalmGRET cells. Cells were imaged by epifluorescence and SRRF microscopy under 488 nm excitation to detect cellular and EP expression of PalmGRET. PalmGRET enables super-resolution imaging of EPs by SRRF microscopy as demonstrated by the comparative micrographs imaged by SRRF (top) and epifluorescence microscopy (bottom) of the boxed regions (i, ii). Individual EPs (arrows) can be resolved in the (i) extracellular space and (ii) cell periphery by SRRF imaging. Epifluorescence microscopy bar, 10 μm; SRRF nanoscopy bar, 1 μm. **(b)** Top, live-cell SRRF imaging of 293T-PalmGRET cells and released EPs. Bar, 1 μm. Bottom, enlarged image of the boxed region exhibiting released EPs. Bar, 200 nm. **(c)** Time-lapse live-cell SRRF imaging of 293T-PalmGRET cells showing an EP protrusion-like event. Bar, 200 nm. See also **Supplementary Movies 2 and 3. (d)** SRRF imaging of lung sections 30 min post-injection of 100 μg EP-293T-PalmGRET or PBS (control). The administered EPs can be detected by SRRF nanoscopy (arrow). The nuclei were stained by DAPI, and EP-293T-PalmGRET was immunoprobed by anti-GFP antibody followed by AlexaFluor 568-secondary antibody. Enlarged images (dashed boxes) of the boxed regions are placed in the top right corner. Bar, 10 μm.

### Visualization, tracking and quantification of EP-PalmGRET from whole animal to nanoscopic resolutions

While our previous GlucB reporter allows *in vivo* imaging of EVs, the signal half-life is short (< 5 min) due to the flash-type emission of *Gaussia* luciferase^[21, 42]^. In addition to facilitating multi-resolution imaging, PalmGRET was designed to enable EP imaging for an extended period of time by employing Nluc, which has been reported to carry a BLI signal half-life greater than 2 h^[43]^. To examine *in vivo* imaging using PalmGRET, 100 μg PBS-washed EP-293T-PalmGRET was IV-injected to C3H mice followed by Fz administration to image the EVs *in vivo* (**Figure 5a**). Of note, 1.2 μm-filtered conditioned medium was used to harvest PalmGRET-labeled EPs to observe the distribution dynamics of multiple EV subtypes and exomeres *in vivo*. We longitudinally tracked *in vivo* the EP distribution signal for up to ~20 min under three different imaging modes (BLI, BRET-FL and FL). Only one brightfield image was acquired at each time point to maximize signal captures under BLI and BRET-FL modes, and hence the signals may not completely overlap with the animal as it gradually shifted to its left over the course of the imaging session. Regions corresponding to the lungs and liver appeared to show the highest signals in the BLI and BRET-FL channels (**Figure 5b**). To confirm this observation, major organs were harvested from EP-injected mice at 30 min and immersed in Fz solution for *ex vivo* imaging. The lungs showed the highest signal followed by the spleen and livers, whereas the brain, heart and kidneys exhibited negligible signals, corresponding to the EP signals detected *in vivo* (**Figure 5c**). While both BLI and BRET-FL channels yielded clear and distinguishable EP signals, the FL channel resulted in a high and non-specific background under external excitation despite its shorter exposure time of 1 s (compared with 60 and 30 s for live animals and harvested organs, respectively, for BLI and BRET-FL).

**Figure 5.**
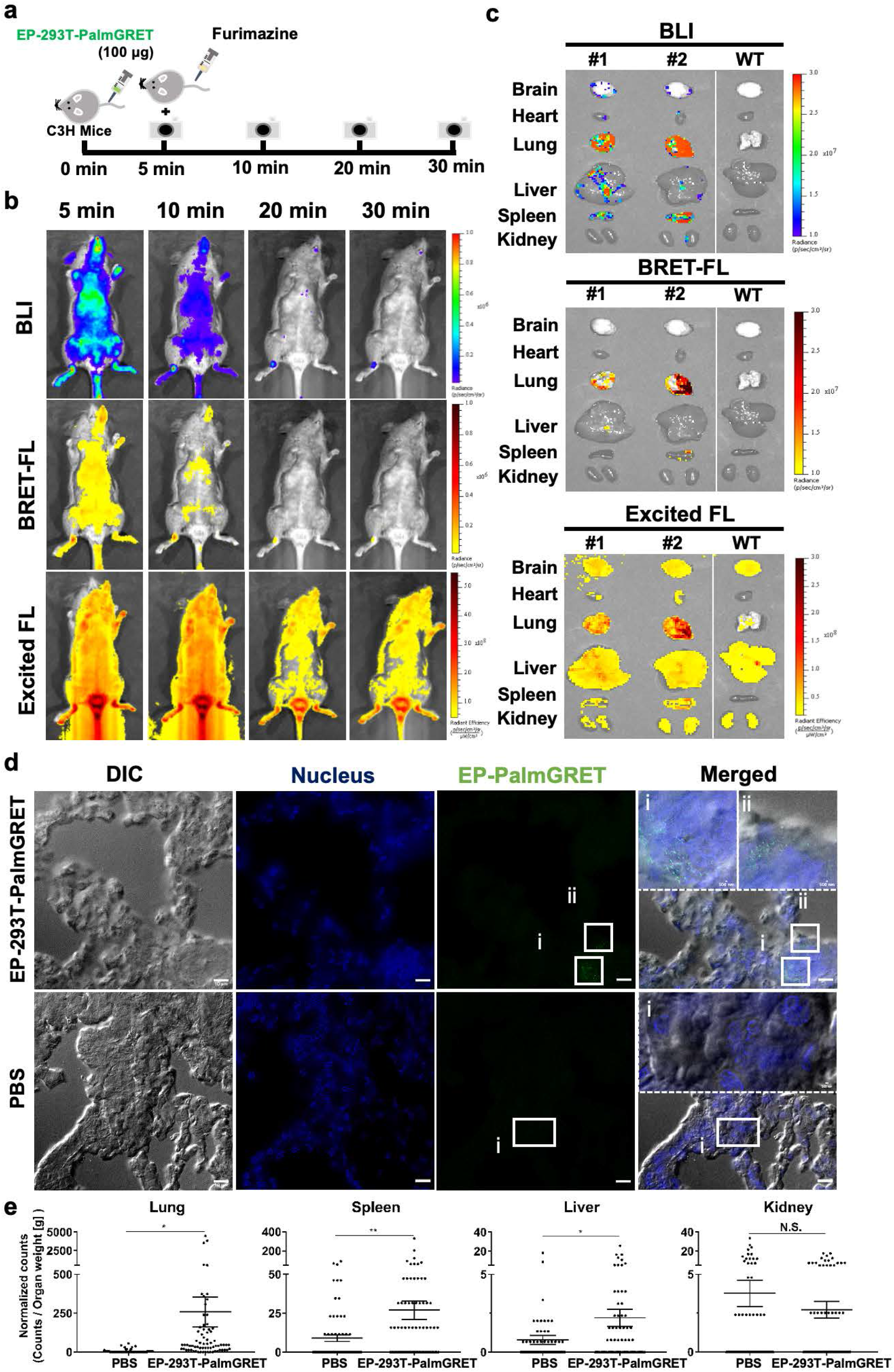
PalmGRET enables multimodal, multi-resolution imaging and analysis of EPs’ distribution in organs. **(a)** Schematic of *in vivo* time-lapsed imaging of EPs. Immunocompetent C3H mice were administered with EP-293T-PalmGRET (100 μg) into the tail vein followed by Fz injection for *in vivo* imaging at 5, 10, 20 and 30 min post-EP administration. **(b)** Bolus IV-injected EP-293T-PalmGRET (100 μg) was visualized under BLI and BRET-FL (GFP using BRET) channels from 5 to 20 min post-administration. The majority of the EP signals were detected at the lungs and spleen. By contrast, imaging under epi-illumination resulted in mostly scattered, non-EP-specific signals. **(c)** *Ex vivo* imaging of organs harvested at 30 min post-injection of EP-293T-PalmGRET (100 μg) (Mice Nos. 1 and 2) and WT subjects. EP-293T-PalmGRET signals were readily detected in the lungs, liver and spleen. **(d)** SRRF nanoscopy of lung sections at 30 min post-injection of EP-293T-PalmGRET (100 μg) or PBS (control). Enlarged images (dashed boxes) of boxed regions (i, ii) reveal injected EPs in lung tissues. The nuclei were stained by DAPI, and EP-293T-PalmGRET was immunoprobed by anti-GFP antibody followed by AlexaFluor 568-secondary antibody to minimize the background signal. Bar, 10 μm; in enlarged images, 500 nm. **(e)** Quantification of EP signals by SRRF imaging demonstrated a significant increase in EP counts in the lungs followed by the spleen and liver at 30 min post-EP injection. The kidneys showed no significant increase in EP counts. 293T-PalmGRET signals were quantified by ImageJ from seventy images of tissue sections of mice injected with EP-293T-PalmGRET or PBS. The EP counts were normalized against organ weight. N.S., *p* > 0.05; *, *p* < 0.05; **, *p* < 0.01 with two-tailed Student’s t-test.

We further visualized EP distributions in the lungs, spleen, liver and kidneys using nanoscopy. The EP-293T-PalmGRET-injected group showed a non-uniform and sporadic distribution of EPs in the lungs (**Figure 5d**). A similar distribution trend was also observed in the spleen, liver and kidneys, suggesting EP uptake by organs is tissue/cell-specific (**Figure S6**). This observation corroborates the findings of Hoshino *et al.* that EVs derived from malignant breast cancer cells are taken up by specific cell types including SPC^+^ epithelial cells and S100A4^+^ fibroblasts^[17]^. Quantification of EP signals from seventy SRRF images per organ with ImageJ^[44]^ showed that the lungs contained the highest EP signals followed by the spleen, liver and kidneys (**Figure 5e** and **Figures S6 and S7**), hence corroborating the EV distribution pattern observed by *in vivo* and *ex vivo* imaging (**Figure 5b, c**). The same imaging and analysis parameters were applied to all tissue samples. Taken together, PalmGRET accurately revealed EP biodistributions with quantification at multiple levels of organization from whole animals and organs to tissues and cells.

### Redirected organotropisms of EPs yield distinct biodistribution profiles

Tumor EVs have been reported to be tropic to pre-metastatic niches to promote subsequent tumor metastasis^[13]^. Other works have decorated EVs with cell type- and cancer-directing peptides such as rabies viral glycoprotein RVG and GE11, respectively, for targeted delivery of therapeutic EVs^[45]^. However, the spatiotemporal dynamics of EV biodistribution following redirection of EV tropism remains largely unknown. To explore this phenomenon, we applied the EP-PalmGRET system to elucidate the spatiotemporal properties of EP in the context of redirected EP tropism. We established the PalmGRET system in a mouse hepatocellular carcinoma cell line, HCA1, which exhibits spontaneous lung metastasis in immunocompetent C3H mice^[46]^, thereby generating stable HCA1-PalmGRET cells. The lung metastatic potential of HCA1-PalmGRET cells was confirmed by orthotopic implantation and detection of tumor formation in the lungs (**Figure 6a, b**).

**Figure 6.**
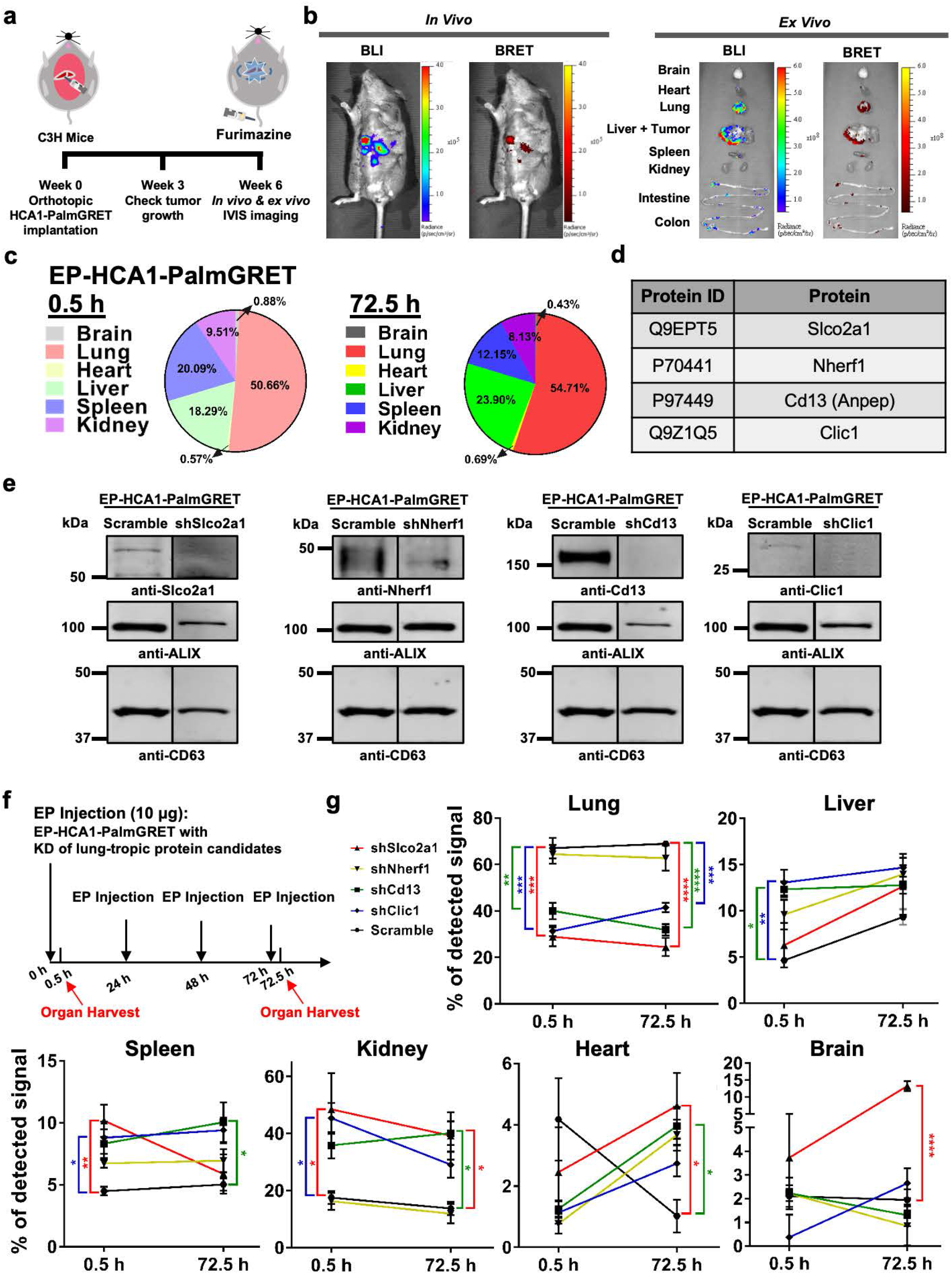
Lung-tropic redirection of EPs yields distinct organ distribution profiles. **(a)** Schematic for tracking spontaneous lung metastasis of orthotopically implanted hepatocellular carcinoma HCA1 in C3H immunocompetent mice. **(b)** Left, *in vivo* imaging under BLI and BRET-FL to visualize liver tumor progression. Right, *ex vivo* imaging of collected organs showing tumors in the livers, as well as lung metastasis. Very low levels of signals were detected at the spleen and intestine. **(c)** Pie charts demonstrating EP-HCA1-PalmGRET predominately localized to the lungs and liver. C3H mice were IV administered with EP-HCA1-PalmGRET (30 μg) at 0, 24, 48 and 72 h. The organs were harvested at 0.5 and 72.5 h for organ distribution analysis. Proportions for each organ at different time points were calculated as 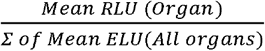. *N* = 3 mice per group with technical triplicates. **(d)** List of lung-tropic EP protein candidates from proteomic analysis of EP-HCA1-PalmGRET. **(e)** Western blot showing knockdown of lung-tropic protein candidates in EP-HCA1-PalmGRET. HCA1-PalmGRET cells were stably transduced with lentivirus encoding shRNA which targets null sequence (Scramble) or individual lung-tropic protein candidates followed by EP isolation and western blot analysis. **Slco2a1**, solute carrier organic anion transporter family member 2A1 (65 kDa); **Nherf1**, sodium-hydrogen antiporter 3 regulator 1 (40 kDa); **Cd13 (Anpep)**, alanine aminopeptidase (110 kDa); **Clic1**, chloride intracellular channel (27 kDa); **β-actin** (42 kDa) as loading control; **Alix**, ALG-2-interacting protein X (95 kDa) and **Cd63**(40kDa) as EV markers. **(f)** Schematic for lung-tropic redirection of EP-HCA1-PalmGRET. C3H mice were IV injected with the EPs (10 μg) at 0, 24, 48 and 72 h through the tail vein, and organs were collected for organ distribution analysis at 0.5 and 72.5 h. **(g)** shSlco2a1, shCd13, and shClic1 significantly reduced lung tropism of EP-HCA1-PalmGRET when compared with the Scramble control. Meanwhile, shSlco2a1 increased EP distribution to the kidney, heart and brain, whereas shCd13 elevated EP distribution to the kidney, heart and spleen at 72.5h. Proportions for each organ at different time points were acquired by 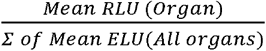. *, *p* < 0.05; **, *p* < 0.01; ***, *p* < L of Mean ELU(All organs) 0.001; ****, *p* < 0.0001 with one-way ANOVA followed by Dunnett’s *post hoc* test compared with the Scramble control. *N* = 3 mice per group with technical triplicates.

As tumor EVs have been shown to target pre-metastatic niches^[13]^, we hypothesized that Eps released by HCA1-PalmGRET will be lung tropic to promote subsequent metastasis. To examine this possibility, we educated immunocompetent C3H mice with 30 μg EP-HCA1-PalmGRET (or PBS as a control; **Figure S8**), and conducted BLI analysis to monitor EP biodistribution in the major organs. Remarkably, EP-HCA1-PalmGRET showed a greater distribution to the lungs and liver (50.7% ± 5.40% and 18.3% ± 3.24%, respectively) at 0.5 h post-injection (**Figure 6c and Figure S9**), indicating its immediate lung-tropic effect. After 72 h, the percentage EP-HCA1-PalmGRET distribution in the lungs remained similar (54.7% ± 3.78%), thereby revealing sustained lung tropism. Meanwhile, the kidneys, heart and brain showed lower percentages of EP distribution (< 10%). Note that 30 μg EP-HCA1-PalmGRET was administered every 24 h based on the finding that this amount yielded minimal signals in the blood circulation 30 min post-injection, suggesting that the majority of the sEVs had been distributed to the organs while leaving a minimal amount in the circulation (**Figure S10a**). While most of the signals were detected in the sEV-enriched qEVoriginal fraction (F8) from the plasma, the neighbouring fractions (F7 and F9) showed lower signals, demonstrating the sEV-labelling specificity of PalmGRET (**Figure S10b**). As C3H mice are immunocompetent, we also performed multiplex cytokine tests on mice serum, and confirmed that the EP injections did not induce an acute immune response, which may have led to a biased uptake in the lungs, livers or spleen (**Table S3**)^[47]^. In addition, since hepatocytes release lipoproteins including palmitoylated proteins ^[48, 49]^, PalmGRET may also label HCA1-PalmGRET-released lipoproteins and contribute to the observed biodistribution profiles. This warrants further investigation.

Given that EP-HCA1-PalmGRET exhibited lung tropism, we next redirected the EPs from the lungs, and investigated the subsequent distribution dynamics to non-targeted organs. We identified several lung tropic protein candidates by proteomic analysis of EP-HCA1-PalmGRET. Among the 521 detected proteins (**Table S4**), we screened for lung tropism-related membrane proteins using Uniprot^[50]^ followed by association to HCC progression, lung metastasis and lung cancer progression, and selected four candidates: Slco2a1^[51]^, Cd13^[52]^, Clic1^[53]^, and Nherf1^[54]^. Following knockdown of the proteins in HCA1-PalmGRET cells (**Figures S11, S12**) and hence reduced expression of the derived EPs (**Figure 6e**), we administered the EPs at 10 μg per 24 h for 72 h to investigate the tropism and biodistribution in C3H immunocompetent mice (**Figure 6f**). At 0.5 h post-injection, the shSlco2a1, shCd13 and shClic1 groups showed a significantly reduced percentages of EPs in the lungs compared with the Scramble control (**Figure 6g, Figure S13**). Interestingly, shCd13 and shClic1 also exhibited increased EPs to the liver, whereas shClic1 and shSlco2a1 demonstrated elevated distribution to the spleen and kidneys at 0.5 h. At 72.5 h, all three groups maintained decreased EP distribution to the lungs. While shCd13 showed increased EP distribution to the spleen, kidneys and heart, shSlco2a1 displayed elevated distribution to the kidneys, heart and brain at 72.5 h. The shNherf1 group yielded neither a change in lung tropism nor an altered biodistribution profile when compared with the Scramble control. A high mRNA level and moderate-high expression of SLCO2A1 have previously been identified in primary and metastatic liver cancers^[51]^, and SLCO2A1 upregulation has been reported to promote lung cancer invasion using a PI3K/AKT/mTOR pathway^[55]^. Downregulation of CLIC1 has been shown to decrease the invasion and migration of HCC cells^[56]^, whereas CLIC1 overexpression in HCC correlates to larger tumor size, distal metastasis, aggressive pathologic tumor phenotype, nodes and poor survival rate^[57]^. Increased CD13 expression has been reported to relate to TGF-β-induced EMT-like phenomenon in liver cancer ^[58]^. Therefore, the loss of lung tropism of EP-HCA1-PalmGRET by Slco2a1, Clic1 and Cd13 knockdowns suggests these proteins direct EP-HCA1-PalmGRET to the lungs for the subsequent lung metastasis of the HCA1-PalmGRET tumor, and warrants further investigation. Additional study will also be required to elucidate the specific tissues and cells facilitating EV uptake by each organ. In summary, we demonstrate that while redirected EP tropism reduces delivery to the lungs, its distribution to other major organs is also dynamically altered, possibly yielding a different function at organ systems level.

## Conclusions

Accurate visualization and tracking of EV distributions are critical to understanding the role of EV-mediated cell-to-cell communication in diseases and therapy. Here we established a multimodal and multi-resolution PalmBRET method to enable pan-EP labelling and imaging and therefore quantification in live cells, whole animals, and preserved tissues. The method can resolve the intricate spatiotemporal dynamics of EPs. PalmGRET revealed that EPs derived from lung metastatic HCC are lung tropic, and the tropism can be mitigated by gene knockdown of the identified lung-tropic membrane proteins. Importantly, the reduced EV delivery to the lungs also significantly alters its distribution to other major organs. Our findings suggest that the dynamics of EV biodistribution and targeted design should be investigated at organ systems level in EP biology and therapeutic developments, respectively.

## Experimental Section

### Cell culture

Human embryonic kidney 293T (HEK293T) cells (a gift from Dr. Chen-Wen Jeff Chan, National Tsing-Hua University) and mouse hepatocellular carcinoma cells from C3H strain HCA1 (a gift from Dan Duda, Massachusetts General Hospital) were maintained in DMEM (high glucose [4,500 mg/L], Hyclone, South Logan, Utah, USA) supplemented with 10% fetal bovine serum (Hyclone), penicillin (100 U/mL) and streptomycin (100 μg/mL) (Hyclone). For stable HEK293T-GRET/PalmGRET cell line maintenance, 1 μg/mL of puromycin (MDbio, Taipei, Taiwan) was added to the culturing medium. Stable HCA1-PalmGRET cells were stored in early passages, and maintained in the culturing medium without puromycin, which was found to affect cellular phenotype. All culturing media were supplemented with plasmocin (2.5 μg/mL; Invivogen, San Diego, California, USA) to mitigate mycoplasma contamination. Both puromycin and plasmocin were removed for two cell passages prior to subsequent experiments.

### Animal studies

C3H/HeNCrNarl male mice were purchased from BioLASCO Taiwan Co., Ltd. (Ilan, Taiwan) for all implantation, EP injection, and EP-tropism studies. For euthanasia, 200 μL Zoletil 50 (25 mg/mL) and xylazine (20 mg/mL) were intramuscularly injected into mice thigh followed by cervical relocation. All animals received humane care in compliance with the Guide for the Care and Use of Laboratory Animals published by the National Academy of Sciences, and all study procedures and protocols were approved by the Animal Research Committee of National Tsinghua University (Hsinchu, Taiwan).

### Molecular cloning

All primers are listed in **Table S1**. pENTR-backbone was created by treating pENTR-Luc (w158-1, a gift from Drs. Eric Campeau and Paul Kaufman; Addgene plasmid #17473) with NcoI and XbaI (New England Biolabs, Massachusetts, USA) followed by gel purification. GRET and PalmGRET fragments were PCR amplified with Q5^®^ High-Fidelity DNA polymerase (New England Biolabs) from pRetroX-Tight-MCS_PGK-GpNLuc (a gift from Antonio Amelio; Addgene plasmid #70185)^[23]^. GRET fragment was flanked by two restriction enzyme sites, NcoI and XbaI at the 5’ and 3’ ends, respectively. The NcoI-GRET-XbaI fragment was then enzyme digested by NcoI and XbaI, gel purified and ligated with pENTR (w158-1)-backbone by Quick Ligation kit (New England Biolabs) to create pENTR-GRET. Palmitoylation signal sequence was added to 5’ end of the GRET fragment through primer synthesis to generate PalmGRET fragment. pENTR-PalmGRET was then constructed *via* NEBuilder HiFi DNA Assembly (New England Biolabs) to insert PalmGRET fragment into the pENTR-backbone. pLenti DEST CMV Puro-GRET/PalmGRET plasmids were created by Gateway LR cloning with pENTR-GRET/PalmGRET and pLenti DEST CMV Puro plasmids (a gift from Drs. Eric Campeau and Paul Kaufman; Addgene plasmid #17452)^[59]^ using LR Clonase II Plus (Invitrogen, Carlsbad, California, USA).

### EP isolation

Stable 293T-GRET/-PalmGRET and HCA1-PalmGRET cells (4.5 × 10^6^ of 293T stable cells or 2.5 × 10^6^ of HCA1-PalmGRET) were cultured in DMEM high glucose supplemented with 5% EV-depleted FBS, penicillin (100 U/mL) and streptomycin (100 μg/mL) for 48 h at 37 °C with 5% CO_2_ in a humidified incubator. Conditioned medium (CM) from cell culture dishes with 80-90% cell confluency were pooled and subjected to differential centrifugation. CM was sequentially centrifuged at 300 × *g* and 2,000 × *g* at 4 °C for 10 minutes to remove cells and cell debris, respectively. CM was then filtered through a 0.22 (for characterization studies), 0.8 (for characterization studies) or 1.2 μm (for characterization studies and tropism studies) pore size PES filter (Pall) and centrifuged at 100,000 × *g* at 4 °C for 90 minutes to collect sEV pellets in double 0.22 μm-filtered PBS.

For tropism study and time-lapse *in vivo* imaging, EPs harvested from 1.2 μm-filtered CM were resuspended with 3 mL double 0.22 μm-filtered PBS and centrifuged under 100,000 × *g* at 4 °C for 70 mins to remove non-EV contaminants. Washed EPs were resuspended in double filtered PBS, and protein concentration was determined by Pierce BCA Kit (Thermo Scientific, Massachusetts, USA). EV samples were aliquoted and stored at −80 °C to maintain same freeze and thaw cycle.

For EP blood clearance study, 30 μg of EP-HCA1-PalmGRET were IV-injected into C3H mice *via* the tail vein. At 0, 5, 30, and 60 min post-injection, the mice were anesthetized by intramuscular injection of 200 μL Zoletil 50 (25 mg/mL) and xylazine (20 mg/mL) cocktail, and blood samples was collected through cardiac puncture. Syringe and sterilized 1.5 mL microcentrifuge tube (Biomate, Taiwan) were pre-rinse by 0.5 M, pH 8 ethylenediaminetetraacetic acid (EDTA, Invitrogen) to prevent blood coagulation. 500-700 μL of mice whole blood was centrifuged at 1,500 *x g* for 10 min at 4 °C, and the supernatant was transferred to a new 1.5 mL microcentrifuge tube for additional centrifugation at 2,000 × *g* for 15 min at 4 °C. The isolated plasma samples were stored at −80 °C prior to EP isolation. For EP isolation from mice plasma, less than 500 μL of mice plasma was loaded into qEVOriginal 70nm columns (iZON Science, Christchurch, New Zealand) based on the manufacturer’s instruction. Double 0.22 μm-filtered PBS was used as the running phase. Fractions 7-9 were individually collected (500 μL/fraction) and concentrated by Amicon Ultra-0.5 centrifugal filter unit with 10 kDa membrane (Merck Millipore) with 14,000 *x g* at 4 °C for 8 min centrifugation to have final volume less than 100 μL. Final volume of the concentrates was recorded for data analysis.

For comparison between sEVs and m/lEVs, CM from 293T-GRET or 293T-PalmGRET were sequentially centrifuged at 300 × *g* and 2,000 × *g* at 4 °C for 10 minutes to remove cells and cell debris, respectively. The supernatants were centrifuged at 10,000 × *g*, 4°C for 30 min to collect 10k pellets (*e.g.* microvesicles; m/lEVs), and the remaining supernatants were further centrifuged at 100,000 × *g*, 4°C for 90 min to harvest 100k pellets (*e.g.* exosomes; sEVs)^[39, 60]^.

### Nanoparticle tracking analysis

NanoSight NS300 (Malvern Panalytical, United Kingdom) was used to determine the size and particle concentration of isolated EVs. NTA Version: NTA 3.2 Dev Build 3.2.16. Triplicate sample measurements using capture camera level: 14 with temperature control of 25 °C ± 0.01°C.

### Transmission electron microscopy and immunogold staining

Isolated sEVs were pelleted at 20,000 x *g* for 30 min at 4 °C followed by fixation with 4% formaldehyde in PBS for 2 h. Fixed sEVs were cryosectioned and immunolabeled with anti-GFP (rabbit; ab6556, Abcam, Cambridge, UK) and anti-CD63 (mouse; 556019, BD Pharmingen, San Diego, USA) followed by rabbit anti-mouse conjugated with 5 nm gold-labeled secondary antibodies (University Medical Center, Utrecht, The Netherlands). Images were captured using Tecnai G^2^ Spirit Bio TWIN transmission electron microscope.

### Sucrose density gradient analysis

For sucrose gradient fractionation, 8, 30, 45, 60 % (w/v) sucrose (J.T.Baker, Pennsylvania, USA) in PBS, pH 7.4 were layered, and isolated sEVs were loaded on top of the discontinuous sucrose gradient followed by centrifugation at 268,000 x *g,* 4 °C for 38 min with MLA-50 rotor (Beckman Coulter, California, USA). The top layer and the subsequent 10 fractions (fraction 1-10) were collected. 16.5 μL of each fraction was saved for BLI activity assay. To pellet sEVs, fractions 2-9 were diluted 1:10 in PBS and centrifuged at 100,000 × *g,* 4°C for 1 h with MLS-55 rotor (Beckman Coulter). The pellets were lysed in 40 μL of radioimmunoprecipitation (RIPA) buffer supplemented with protease inhibitor cOmplete protease inhibitor cocktail (Roche, Basel, Switzerland). 6.6 μL of each pelleted fraction was saved for BLI activity assay.

### Western blot analysis

Cells and EPs were lysed in RIPA lysis buffer (1X PBS, pH 7.4, with 1% Igepal CA-630, 0.5% sodium deoxycholate, 0.1% SDS) with cOmplete protease inhibitor cocktail (Roche). Protein concentrations of EPs and cell lysates were determined using Pierce™ BCA Protein Assay Kit (Thermo Scientific). Protein lysates were separated using Bolt™ 4-12% Bis-Tris Plus Gels (Invitrogen) and transferred onto 0.2 μm nitrocellulose membranes (Amersham Bioscience, Little Chalfont, UK). The membranes were blocked with 5% BSA (Cyrusbioscience, Taiwan) in PBST (1X PBS, pH 7.4, with 0.1% Tween-20) for 1 h at room temperature and immunoprobed with primary antibodies overnight at 4 °C. The membranes were washed three times in PBST at room temperature, incubated with secondary antibodies for 1 h at room temperature followed by another three washes in PBST. For Western blots of purified sEV pellets from sucrose gradients, the membranes probed by HRP-conjugated secondary antibodies were developed by chemiluminescence using Trident Femto Western HRP Substrate kit (GeneTex) with MultiGel-21 Imaging System (Topbio, Taiwan). For western blots of protein knockdown, EV-293T-GRET/-PalmGRET subtypes, and EP-HCA1-WT/GRET/PalmGRET characterization experiments, the targeted proteins immunoprobed by IR-dye-conjugated secondary antibodies were imaged by ODYSSEY CLx (LI-COR Biosciences, Nebraska, USA). Antibody dilutions and hosts are provided in **Table S2**.

### Dot blot analysis

To characterize membrane orientation of PalmGRET and GRET in sEVs, sEVs isolated from 0.22 μm-filtered CM were serially diluted to the following concentrations: 31.25 ng, 62.5 ng, 125 ng, 250 ng, 500 ng, 1,000 ng, 2,000 ng in 5 μL. Cell lysates were diluted to 2,000 ng in 5 μL as positive controls. 5 μL of diluted sEVs, cell lysates, and double-filtered PBS were dotted onto 0.45 μm nitrocellulose membranes (Amersham Bioscience) and blocked in 10% BSA supplemented PBS overnight at 4 °C. The membranes were immunoprobed with anti-GFP antibody (GeneTex) diluted in 5% BSA in PBS or PBST overnight at 4 °C, wash three times in PBS or PBST for 30 minute each, and incubated with HRP-conjugated secondary antibodies for 1 h at room temperature followed by another 3 washes. The membranes were developed by chemiluminescence using ECL Select™ Western Blotting Detection Reagent (Amersham Bioscience) and MultiGel-21 Imaging System (Topbio). Antibody dilutions and hosts are provided in **Table S2**.

### Asymmetric-flow field-flow fractionation (AF4)

AF4 of sEV-GRET/-PalmGRET was performed based on the protocol described by Zhang and Lyden ^[61]^ with minor modifications. Briefly, 4.5 × 10^6^ 293T-PalmGRET or -GRET cells were seeded in 15 cm culture dishes (Thermo Scientific) with DMEM high glucose (Hyclone) supplemented with 5% EV-depleted FBS, penicillin (100 U/mL, Hyclone) and streptomycin (100 μg/mL, Hyclone) for 48 h at 37 °C with 5% CO_2_ in a humidified incubator. After 48 h, CM were pooled into 50 mL centrifuge tubes (Biomate) and centrifuged at 500 × *g* for 10 min, 10°C followed by 12,000 × *g* for 20 min, 10°C. CM were then centrifuged at 100,000 × *g* for 90 minutes, 4 °C with 70Ti rotor (Beckman Coulter) to collect sEV pellets in double 0.22 μm-filtered PBS. The harvested sEVs were resuspended with 3 mL double filtered PBS and centrifuged with MLA-55 rotor (Beckman Coulter) under 100,000 × *g* at 4 °C for 70 mins to remove non-EV contaminants. The washed samples were resuspended in double filtered PBS, and protein concentration was determined by Pierce BCA Kit (Thermo Scientific). sEV samples were adjusted to 0.5 mg/mL and stored at −80 °C. Before loading to AF4, the frozen sEV-GRET and - PalmGRET were thawed on ice and centrifuged at 13,000 × g at 4°C for 5 min. 100 μL of the supernatant was injected to AF4 (Eclipse DualTec; Wyatt, California, USA) to collect the sEV subpopulations. The same AF4 parameters were set as described in Zhang ****et al.****^[61]^. Resulting fractions were collected in 1 mL deep 96-well plate (Greiner Bio-One, Germany) for 500 μL/well. P2, P3 and P4 comprise of fractions from 13.5 min to 19 min, 22 min to 27 min, and 31 min to 34.5 min, respectively. The pooled fractions were transferred to ice-cold PBS pre-rinsed Amicon Ultra-4 centrifugal filter units with Ultracel-30 membrane (Merck Millipore, Massachusetts, USA), then centrifuged with 3,200 *x g* at 4 °C for 10 min. The centrifugation was repeated until the concentrate volume was less than 100 μL. The final volume of each population was recorded for data analysis.

### Live-cell imaging

To setup live-cell fluorescent imaging, 1 × 10^4^ 293T-GRET/-PalmGRET cells were seeded per well into a glass-bottom 8-well chamber (#1.5, Nunc™ Lab-Tek™ II Chamber Slide™ System, Thermo Scientific) coated with poly-D-lysine (Mw 75,000-150,000 Da, Sigma-Aldrich), and incubated overnight in 250 μL DMEM high glucose without phenol red (Hyclone) supplemented with 10% FBS (Hyclone), penicillin (100 units/ml, HyClone) and streptomycin (100 μg/ml, HyClone). Cells were then imaged at 37 °C in a chamber supplemented with humidified 5% CO_2_ through 63x oil lens (NA= 1.4) with optovar set to 1.6x using Axio Observer 7 epifluorescent microscope (Zeiss, Oberkochen, Germany) and recorded by iXon Ultra 888 EMCCD (Andor Technology, Northern Ireland, UK). Micro-Manager ^[62]^ was used to acquire the images with or without SRRF ^[35]^, and all images were taken with the same parameter for DIC, GFP (epi-FL), or GFP (BRET), or GFP (SRRF-FL). Images of GFP *via* BRET were acquired after adding furimazine (Promega, Wisconsin, USA) diluted in culturing medium without phenol red (final dilution: 1/500) into an 8-well chamber containing either 293T-GRET/PalmGRET cells, then imaged by applying 530/50 nm emission filter without excitation light. To setup live-cell confocal imaging for Z-stack reconstruction, 3 × 10^4^ 293T-PalmGRET or 1.5 x 10^4^ for both 293T-PalmGRET and 293T-PalmtdTomato cells were seeded into 35 mm glass-bottom (#1.5) dish (ibidi, Gräfelfing, Germany) coated with poly-D-lysine (Mw 75,000-150,000 Da, Sigma-Aldrich) and incubated overnight in 1.5 mL DMEM high glucose (Hyclone) supplemented with 10% fetal bovine serum (Hyclone), penicillin (100 units/ml, HyClone) and streptomycin (100 μg/ml, HyClone). Cells were then imaged at 37°C in a chamber supplemented with humidified 5% CO_2_ through white light laser confocal microscope TCS SP8 (Leica, Wetzlar, Germany). **Movie S1** was processed by Bitplane-Imaris 9.5 (Andor Technology) from images of **Figure 1e** and **Movie S2 and S3** were processed by Fiji (ImageJ, NIH).

### HCA1-PalmGRET orthotopic implantation in C3H mice

C3H mice (Biolasco) were anesthetized, and 1 × 10^6^ cells of HCA1-PalmGRET were mixed with 10 μL Matrigel (Corning, New York, USA) then orthotopically injected to the liver of 5-6 weeks old C3H mice as previous described ^[46]^.

### Proteomics analysis and data processing

EP samples harvested from 1.2 μm-filtered CM were resolved in 10 mM tris (2-carboxyethyl) phosphine (Sigma-Aldrich) with 50 mM ammonium bicarbonate (Sigma-Aldrich) for 30 min at 37 °C and alkylated in 50 mM iodoacetamide (Sigma-Aldrich) with 50 mM ammonium bicarbonate for 45 min in the dark at 25°C. Trypsin/Lys-C (Promega) solution was added and incubated at 37 °C for 12 h. The resulting peptides were extracted from gel fragments and analyzed with Orbitrap Fusion Lumos Mass Spectrometer (Thermo Scientific) combined with UltiMate 3000 RSLC nano-flow HPLC system (Thermo Scientific) with HCD or EThcD MS/MS mode. Raw file of MS/MS spectra were processed for peak detection by using MaxQuant software version 1.6.5.01.^[63]^ Peptide identification was performed by using the Andromeda search engine^[64]^ and the Swiss-Prot database. EP-HCA1-PalmGRET was searched against UniProt Mus Musculus Database (Swiss-prot, 17,009 reviewed protein sequence, UP000000589). Search criteria used in this study were trypsin specificity, fixed modification of carbamidomethyl, variable modifications of oxidation and acetyl (Protein N-term), and allowed up to 2 missed cleavages. A minimum of 5 amino acids in the peptide length was required. The precursor mass tolerance was 20 ppm, and the fragment ion tolerance was 0.5 Da. False discovery rate of 0.01 and Peptide Rank of 1 were set for peptide identification filters.

### EP-tropism study

30 μg of EP-HCA1-PalmGRET or PBS was IV-injected to C3H mouse *via* the tail vein at 0, 24, 48, and 72 h. At 0.5 and 72.5 h post-EP administration, each mouse was anesthetized and euthanized with cervical dislocation, and major organs including the brain, lungs, heart, livers, spleen, and kidneys were harvested for biodistribution analysis by BLI assay. Total RLU per organ was calculated as follows: (RLU/20 μl) x (organ/mL T-PER lysis buffer). To study redirected EP tropism, pLKO.1 short hairpin ribonucleic acid (shRNA) backbone encoding mouse Slco2a1 (TRCN0000079869), Nherf1 (TRCN0000068587), Clic1 (TRCN0000069738), Cd13 (TRCN0000031088), or Scramble (ASN0000000002) from National RNAi Core Facility (Academia Sinica, Taipei, Taiwan) were individually packaged into lentiviruses as described in the lentivirus production section (**Table S5**). HCA1-PalmGRET cells were then infected with the lentiviruses establish knockdown cell lines in 15 cm dishes. At 24 h post-infection, 2.5 × 10^6^ transfected cells per dish were transferred to 3-4 × 15 cm culture dishes with DMEM high glucose (Hyclone) supplemented with 5% EV-depleted FBS (Hyclone), penicillin (100 U/mL, Hyclone) and streptomycin (100 μg/mL, Hyclone) for 48 h at 37 °C with 5% CO_2_ in a humidified incubator followed by EP isolation. Timelines for EP administration (10 μg of EP injected at each specified time point) and organ harvest are described in **Figure 6**. Proportions of detected EP signal was calculated as follows: (RLU/specific organ) / (RLU/all organs) × 100%.

#### RNA isolation and real-time quantitative polymerase chain reaction (RT-qPCR)

Total RNAs were isolated from HCA1-PalmGRET cells using RNeasy Mini Kit (QIAGEN) and subjected to RT-qPCR to quantify mRNA expressions of: Slco2A1, Cd13, Clic1, Nherf1, β-actin. Total RNA were first reverse transcribed into complementary DNA using SuperScriptTM IV First-Strand Synthesis System (Invitrogen) followed by qPCR with SYBR Green PCR Master Mix (Invitrogen) using a StepOnePlus Real-Time PCR System (Applied Biosystem: 95°C for 10 min, 40 cycles at 95°C for 15s, and 62°C for 1 min. The relative expressions of genes were calculated using ΔΔCT method. Primers are listed in **Table S6**.

### Immunohistochemistry

The kidneys, livers, lungs, and spleen were embedded in Tissue-Tek (OCT compound) and kept frozen at −80 °C. Tissues were sectioned (5 μm thickness), fixed in acetone at −20 °C for 10 min and washed with PBS. The sections were blocked with 5% BSA for 1 h at room temperature and incubated with primary antibodies at 4 °C overnight. Sections were washed with PBS and incubated with secondary antibodies for 1 h at room temperature. Sections were next washed by PBS and mounted using Prolong Diamond antifade mounting solution containing DAPI (Invitrogen). Primary and secondary antibody dilution for different organ tissues are listed in **Table S2.** All sections were imaged using Axio Observer 7 epifluorescent microscope (Zeiss) equipped with iXon Ultra 888 EMCCD (Andor Technology) with or without SRRF with the same imaging parameters. Images (n = 70 per organ) were subjected to Fiji (ImageJ, NIH) for EP quantitative analysis with the same threshold. 5 × 5 tiled images were selected from 6 × 6 tiled images to remove uneven edges following tiling by Fiji (NIH).

### Bioluminescence assays

GloMax® Discover Microplate Reader (Promega) with auto-injectors was set for Nluc (450/10 nm) and BRET-FL (510/10 nm) readings, and BRET ratio was calculated by dividing the acceptor signal (510/10 nm) with the donor signal (450/10 nm). To assess EV BL, 5 μL of crude EVs (triplicates), 5 μL of each fraction (triplicates), or 3 μL of pelleted fraction 2-9 (duplicates) were added into a 96-well white plate (Greiner Bio-One, Germany). Furimazine (Promega) was diluted 1:100 in PBS and 50 μL was added to each well by the auto-injector to detect Nluc and BRET signals with 0.5 sec integration time. To assess AF4 samples, 20 μL concentrates of P2, P3, and P4 of sEV-293T-GRET/PalmGRET (triplicates) were added into a 96-well white plate (Greiner Bio-One). Furimazine (Promega) was diluted 1:400 in PBS and 80 μL was injected into each well by the auto-injector to detect Nluc and BRET-FL signals with 1 sec integration time. To track time-lapse sEV/EP-PalmGRET/GRET signal, isolated sEVs/EPs were re-suspended in double-filtered PBS and 5 μL of sEVs/EPs were added per well (triplicates) into a 96-well white plate (Greiner Bio-One). Furimazine (Promega) was diluted 1:500 in PBS, and 45 μL of Fz solution was injected into sEV/EP-293T-GRET/PalmGRET *via* the auto-injector. BL and BRET-FL signals were measured every 5 minutes for a total of 4 h with 0.5 sec integration time. The collected signals were normalized to the number of cells or sEV/EP particles determined by a cell counter (Countess II™, Invitrogen) or NTA analysis, respectively.

For EP biodistribution analysis, the harvested organs were homogenized and lysed by T-PER (0.5 g organ/mL T-PER, Thermo Scientific) with 1X Halt protease inhibitor (Thermo Scientific). Lysates were centrifuged at 10,000 *x g*, 4°C for 10 minutes followed by supernatant collection to acquire protein extracts from tissues. The supernatants were added to 96-well white plate (Greiner Bio-One) with 20 μL per well in triplicates. 80 μL of furimazine (Promega) with 1/400 dilution in PBS was auto-injected per well to have final dilution of 1/500. Integration time was set to 1 s for BLI readings.

To monitor time-lapse PalmGRET/GRET signals of live cells, 1 × 10^4^ 293T-PalmGRET, - GRET, and -WT cells were seeded per well in triplicates in a 96-well white plate coated with poly-D-lysine (Mw 75,000-150,000 Da, Sigma-Aldrich), and incubated overnight in 100 μL DMEM high glucose without phenol red (Hyclone) supplemented with 10% fetal bovine serum (Hyclone), penicillin (100 units/ml, HyClone) and streptomycin (100 μg/ml, HyClone). 50 μL culture medium was next removed from each well, and 50 μL furimazine (Promega) diluted 1:250 in the culturing medium was auto-injected into each well for detection of bioluminescent and BRET signals every 5 minutes for a total of 8 h with 0.5 sec integration time.

To measure plasma-derived EPs, 20 μL concentrates of fraction 7-9 from mice plasma (by qEV columns) were add into a 96-well white plate in triplicates (Greiner Bio-One). Furimazine (Promega) was diluted 1:400 in PBS, and 80 μL was injected into each well by an auto-injector to detect Nluc and BRET-FL signals with 1 sec integration time.

### *In vivo* and *ex vivo* imaging

To image biodistribution of administered EPs *in vivo*, 100 μg of EP-293T-PalmGRET (PBS washed) was IV-injected to C3H mouse *via* the tail vein at 0 min. The mice were anesthetized at 3 mins post-EP injection. At 5 mins post-EP injection, 1/40 diluted furimazine (Promega) in sterile 100 μL PBS was IV-injected to C3H mice *via* tail vein for IVIS imaging. Images were taken at 5, 10, 20, and 30 min post-Fz administration. Following the initial brightfield imaging, Nluc and BRET-FL signal were acquired by applying open filter (no filter) and 500/20 nm, respectively. For epifluorescence signal, excitation filter (470/30 nm) and emission filter (520/20 nm) were used. 60 and 30 sec exposure time were set for *in vivo* and *ex vivo* images, respectively; and 1 sec was set for externally excited fluorescence images to avoid overexposed images.

Prior to monitoring tumor progression, HCA1-PalmGRET bearing C3H mice were anesthetized. Following the initial brightfield imaging, Nluc and BRET-FL signals were sequentially acquired by IV-injection with 1/40 diluted furimazine (Promega) in 100 μL sterile PBS and applying open (for Nluc) and 500/20 nm (for BRET-FL) emission filters with 60 sec exposure time. Following live animal imaging, each mouse was anesthetized again, and euthanized with cervical dislocation. Major organs including the brain, lungs, heart, livers (with tumor), spleen, kidneys, intestine and colon were harvested for *ex vivo* imaging. The harvested organs were immersed in 1/400 diluted furimazine (Promega) in sterilized PBS to detect Nluc/BRET-FL signal. The same filter settings were applied to collect Nluc and BRET-FL with 1 sec exposure time to avoid saturated signals.

### Multiplex cytokine assay of mouse serum

Mice were anesthetized and the blood was collected by cardiac puncture at 0.5, 24.5, 48.5 and 72.5 h post-EP-HCA1-PalmGRET injection. The blood of mice with PBS injection were included as the baseline (control). 500-700 μL of mice whole blood were collected in 1.5 mL and allowed to clot at 4 °C for 30 min before centrifugation. After centrifugation with 1,500 *x g* at 4 °C for 10 min, the serum were transferred to a new 1.5 mL microcentrifuge and stored at −80 °C before Multiplex Cytokine Assay. Bio-Plex® Multiplex Immunoassays Assay (Bio-Rad, California, USA) was performed according to manufacturer’s instruction. 50 μL serum was loaded per well into 96-well microplate, and each sample was test in duplicates. Murine IL-1β, IL-2, IL-4, IL-5, IL-6, IL-10, IL-12, IL-13, IL-17A, IL-22, IL-23, TNF-α, and IFN-γ signal were test by the Bio-Plex array reader (Bio-Rad).

### Statistical analyses

Statistical analyses were performed using GraphPad Prism Version 7.05. Two-tailed Student’s t-test was used for comparison between two groups. Two-way analysis of variance (ANOVA) followed by Tukey *post hoc* test, or ordinary ANOVA followed by Tukey or Dunnett *post hoc* test was used for comparison of three or more groups. Values were normally distributed, and the variance was similar between compared groups. Error bars for all the graphs represent mean ± SEM. P-value < 0.05 was considered statistically significant.

## Supporting Information

Supporting Information is available from the Wiley Online Library.

## Acknowledgements

We thank Dr. Wei Chun Huang (Biophysics Core Facility, Institute of Atomic and Molecular Sciences) for assistance on confocal laser microscopy. We are grateful to Chia Chen Tai and Tzu Wen Tai [Flow Cytometry Core Facility, Academia Sinica Core Facility and Innovative Instrument Project (AS-CFII108-113)] for assistance on the cell sorting service. We thank Inflammation Core Facility (Institute of Biomedical Sciences, Academia Sinica, Taiwan) for assistance on multiplexed murine cytokine assay. We thank the National RNAi Core Facility (Academia Sinica, Taiwan) for providing the shRNA plasmids. The Lai Lab is supported by the Ministry of Science and Technology (MOST) grants 104-2320-B-007-005-MY2 (C.P.L.), 106-2320-B-007-004-MY3 (C.P.L.), Academia Sinica Innovative Materials and Analysis Technology Exploration (i-MATE) Program AS-iMATE-107-33 (C.P.L.), and Academia Sinica Career Development Award AS-CDA-109-M04 (C.P.L.). A.Y.W. and C.P.L. conceived and designed the study. A.Y.W., J.J.K. and A.L.A. performed initial experiments. A.Y.W., Y.C., V.G., J.C.C., S.W. and W.L. performed characterization experiments. A.Y.W., Y.S., Y.C., H.H. and Y.C. performed animal studies. S.T.C., C.H.Y.C., H.J., K.U. performed and analyzed proteomic study. A.Y.W., J.C. and M.H. performed and analyzed AF4 experiment. M.E. performed and analyzed TEM imaging. A.Y.W., Y.C., V.G., and C.P.L. analyzed data. C.P.L. supervised the study. The manuscript was written by A.Y.W. and C.P.L. with input from all authors.

## Table of contents

PalmGRET, a bioluminescence resonance energy transfer (BRET)-based reporter for extracellular particles (EPs) enables pan-EP labeling, including extracellular vesicles (EVs) and exomeres. PalmGRET allows accurate visualization, tracking and quantification of EPs from whole-animal to nanoscopic resolutions under different imaging modalities, including bioluminescence, BRET and fluorescence. Using PalmGRET, we identified lung tropic EP proteins and revealed dynamically altered biodistributions under redirected tropism.

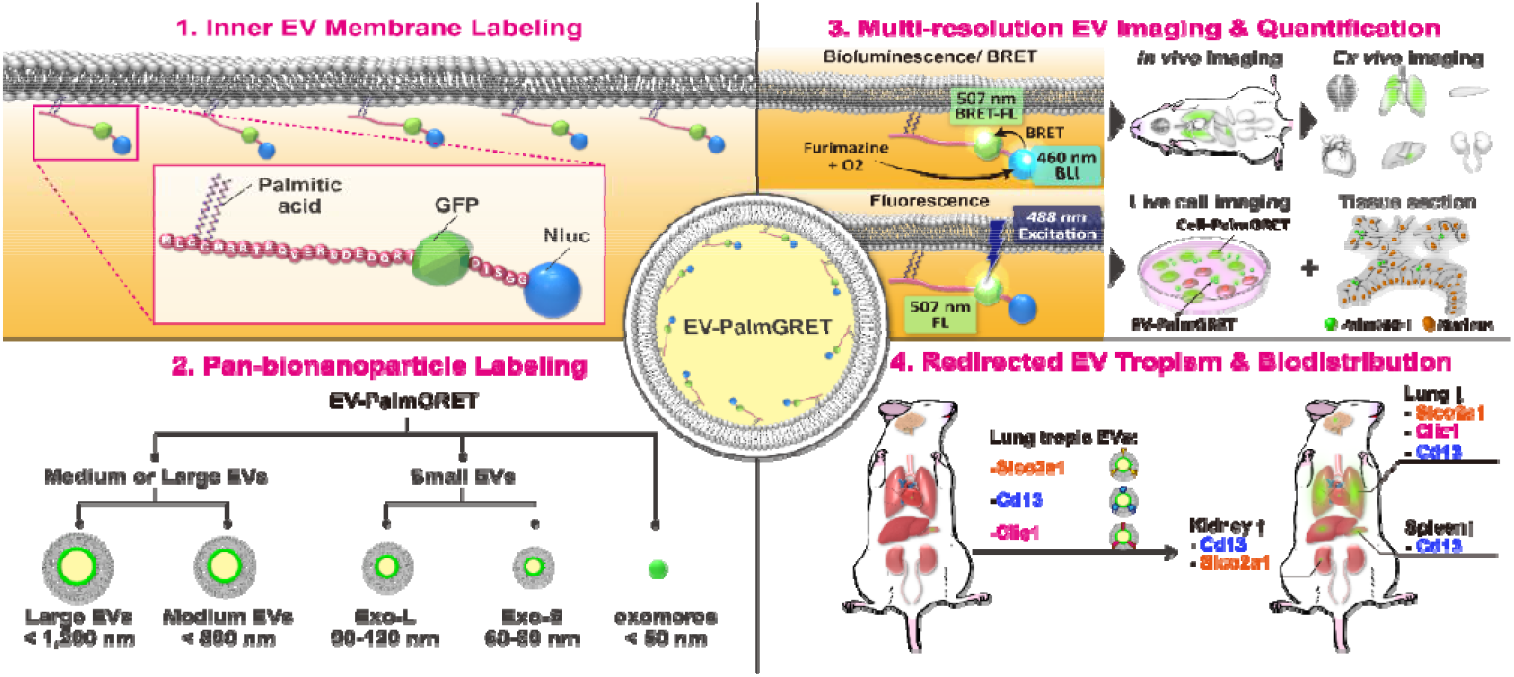

