## Supplementary_Tables_Figures for "Multi-resolution imaging using bioluminescence resonance energy transfer identifies distinct biodistribution profiles of extracellular vesicles and exomeres with redirected tropism"

**Supporting Information**

13 Supplementary Figures

6 Supplementary Tables

3 Supplementary Movies

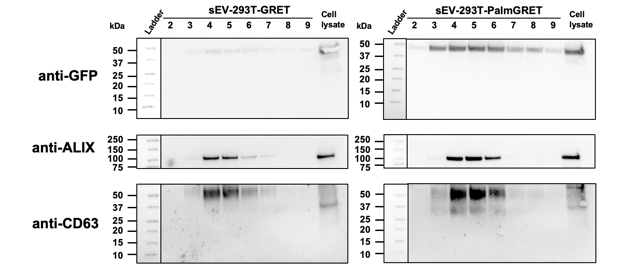

**Figure S1. Original western blots of sEV-GRET and sEV-PalmGRET following sucrose gradient purification.** Nitrocellulose membranes (pore size: 0.45 μm) were cut into two blots to acquire Alix and CD63 signals on the first day. The GFP of GRET/PalmGRET was detected the next day after treating the CD63 blot with stripping buffer and immunoprobing with anti-GFP antibody. Expected size: Alix (95 kDa), CD63 (30–60 kDa), GRET (46 kDa) and PalmGRET (49 kDa)

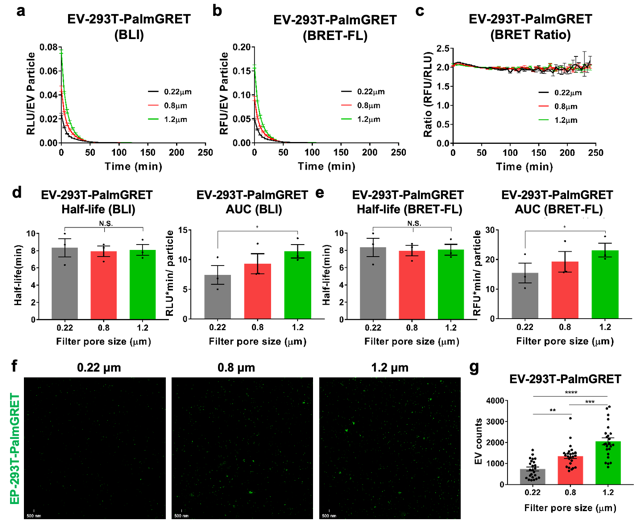

**Figure S2. PalmGRET labels small, medium and large EVs.** Conditioned medium from 293T-PalmGRET cell were filtered with a 0.22, 0.8 or 1.2 μm filter to examine the labelling ability of EV subtypes by PalmGRET. 5 μL EVs from each group was used to record time-lapse data for **(a)** BLI, **(b)** BRET-FL and **(c)** BRET ratio. Representative charts from one of three independent experiments are shown. Half-life and AUC of **(d)** BLI half-lives for the 0.22, 0.8 and 1.2 μm samples are 8.3 ± 1.07, 7.93 ± 0.61 and 8.08 ± 0.62 min, respectively; the BLI AUCs are 7.4 ± 1.58, 9.3 ± 1.69 and 11.4 ± 1.13 RLU·min/particles, respectively. **(e)** BRET-FL half-lives for the 0.22, 0.8 and 1.2 μm samples are 8.3 ± 1.06, 8.0 ± 0.60 and 8.07 ± 0.63 min, respectively; the BRET-FL AUCs are 15.4 ± 3.37, 19.3 ± 3.49 and 23.2 ± 2.35 RFU·min/particles, respectively. N.S., *p* > 0.05; *, p< 0.05 with one-way ANOVA followed by Tukey’s *post hoc* test for three independent experiments. (**f, g**) 3 μL EV was used for SRRF nanoscopy and EV quantification. **(f)** SRRF nanoscopy of EV-293T-PalmGRET isolated from 0.22, 0.8 or 1.2 μm filtered CM. Bar, 500 nm. **(g)** Quantification of EVs from SRRF nanoscopy images demonstrates a positive correlation between EV count and the filter pore sizes. Eight fields of interest were taken from the EVs of three independent experiments (*N* = 24). **, *p* < 0.01; ***, *p* < 0.001; ****, *p* < 0.0001 with one-way ANOVA followed by Tukey’s *post hoc* test.

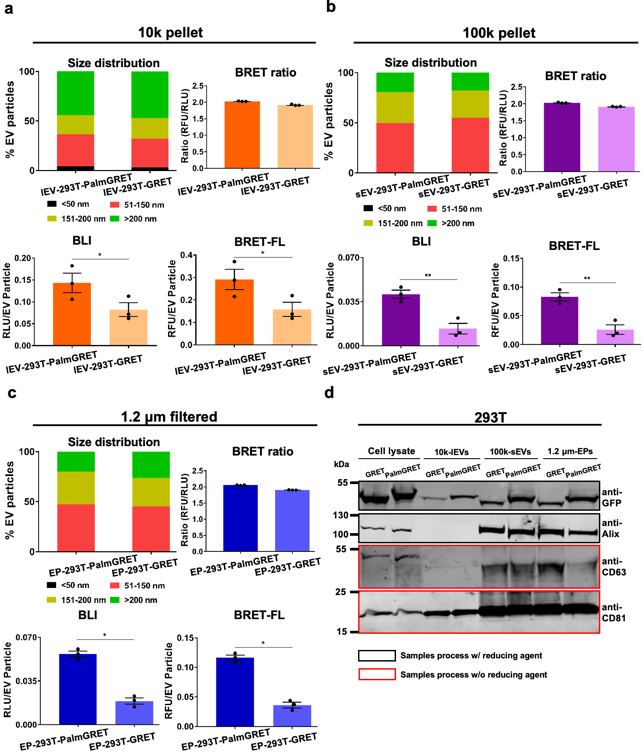

**Figure S3. Characterization of 293T-GRET/-PalmGRET-derived large EVs and small EVs.** NTA, BLI activity, BRET-FL, and BRET ratio of EV pellets harvest from: **(a)** 10k (10,000 × *g*) centrifugation; **(b)** 100k (100,000 × *g*) centrifugation, and; **(c)** 1.2 μm filtered CM followed by 100,000 × *g* centrifugation. 100k and 1.2 μm-filtered samples exhibited a similar size distribution. By contrast, the 10k sample exhibited a size distribution shifted towards the >200 nm fraction. *, *p* < 0.05; **, *p* < 0.01 with two-tailed Student’s t-test. **(d)** Western blot analysis of 10k-m/lEVs, 100k-sEVs, and 1.2 μm filtered-EPs from 293T-GRET/-PalmGRET cells. In 293T-10k-m/lEVs, Alix was not detected and CD63 faintly observed, whereas the CD81 was found less compared to 100k-sEVs, and 1.2 μm filtered-EPs.^[4,39,65]^ Black and red squares indicate the samples were processed with and without 50 mM dithiothreitol (DTT) reducing agent, respectively. Expected size: GRET (46 kDa); PalmGRET (49 kDa); Alix (96 kD); CD63 (26 kDa); CD81 (26 kDa).

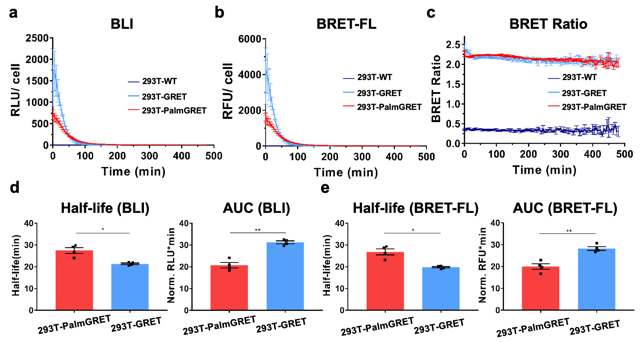

**Figure S4. Nluc and BRET activities of stable 293T-PalmGRET and 293T-GRET cells. (a)** 293T-GRET cells showed a higher Nluc activity from 0 to 60 min following Fz administration than either 293-PalmGRET cells or wildtype 293T cells (293T-WT; control). 293T-WT exhibited a limited amount of Nluc background activity. **(b)** BRET-excited GFP fluorescence was simultaneously collected in the same experiment, and 293T-GRET demonstrated a stronger GFP signal using BRET than 293T-PalmGRET from 0 to 60 min. 293T-WT showed no detectable GFP signal. (**c**) 293T-PalmGRET and 293T-GRET exhibited a significantly higher and similar BRET ratio over time than 293T-WT control. **(d)** Half-life and AUC of Nluc. **(e)** Half-life and AUC of BRET-excited GFP. *, *p* < 0.05; **, *p* < 0.01 with two-tailed Student’s t-test.

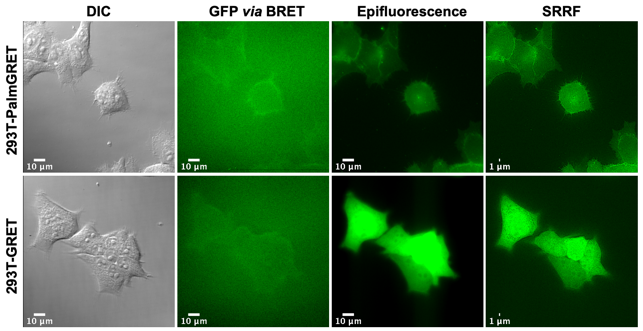

**Figure S5. Live-cell, multimodal imaging of 293T-PalmGRET and 293T-GRET cells.** The cells were treated with Fz and imaged by an EMCCD camera to detect BRET-excited GFP signals. Bar, 10 μm. The same samples were next imaged by epifluorescence (Bar, 10 μm) and SRRF (Bar, 1 μm) microscopy under 488 nm excitation to detect cellular and EV expression of PalmGRET.

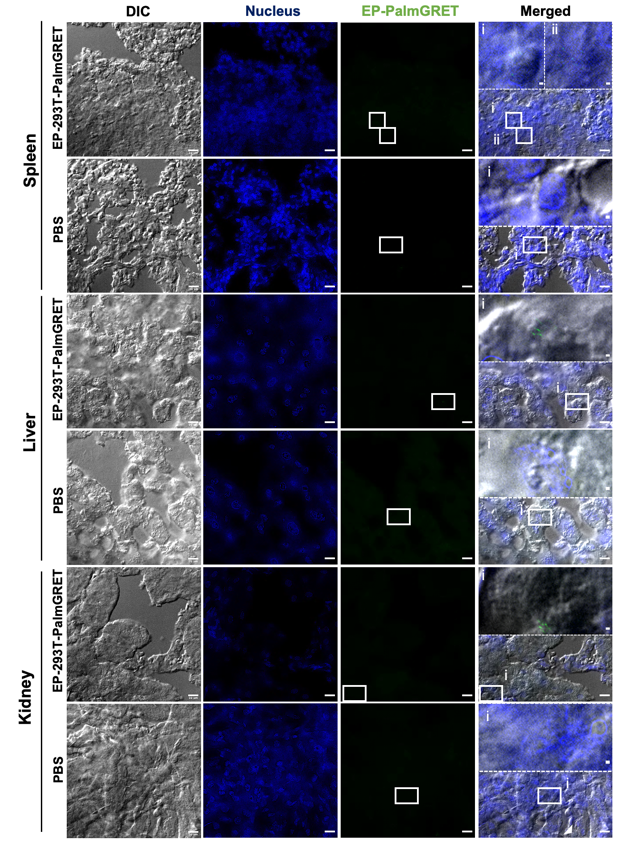

**Figure S6. SRRF nanoscopy of organs from EP-293T-PalmGRET-administered C3H immunocompetent mice.** SRRF and DIC images of the spleen, liver, and kidney sections at 30 min post-EP-293T-PalmGRET (100 μg) or PBS (control) injection. DAPI was used for nucleus staining; primary anti-GFP and Alexa Fluor^®^ 568-conjugated secondary antibodies were used to probe PalmGRET. Bar, 10 μm. Enlarged images (dashed boxes) of boxed region (i, ii) are placed at the top of the merged images. Bar, 500 nm.

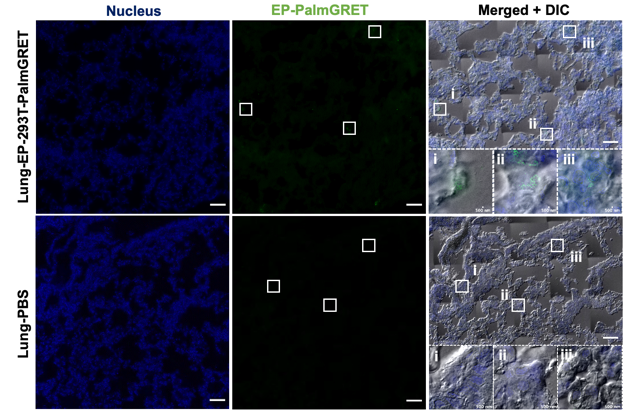

**Figure S7. Mosaic SRRF images reveal an uneven distribution of EP-293T-PalmGRET in the lungs of C3H mice.** 5 × 5 mosaic SRRF and DIC images of C3H mouse lung tissue injected with EP-293T-PalmGRET (top, 100 μg) and PBS (bottom). DAPI was used for nucleus staining; anti-GFP primary antibody followed by Alexa Fluor^®^ 568-conjugated secondary antibody were used for PalmGRET staining. Bar, 50 μm. Enlarged images (dashed boxes) of boxed regions (i–iii) indicate detected EV or background signals (control). Bar, 500 nm.

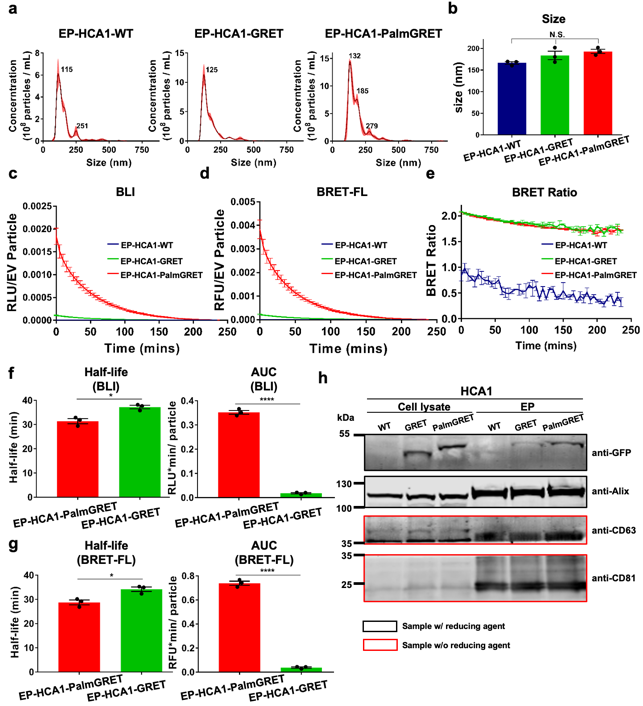

**Figure S8. Characterization on mouse hepatocellular carcinoma HCA1-derived EPs.** EPs were harvested from 1.2 μm-filtered CM of HCA1-WT, -GRET, -PalmGRET cells with 100,000 × *g* centrifugation. **(a, b)** NTA showed a similar size distribution (**a**) and mean size (**b**) between EV-HCA1-WT, -GRET, and -PalmGRET (160.2 ± 5.11, 183.5 ± 16.96, and 193 ± 8.72 nm). N.S., *p* > 0.05 with one-way ANOVA followed by Tukey’s *post hoc* test with three independent experiments. **(c–e)** 5 μL EVs from each group was used to record time-lapse data for **(c)** BLI, **(d)** BRET-FL and **(e)** BRET ratio. Representative charts from one of the three independent experiments are shown. (**f**) BLI half-life and AUC of EP-PalmGRET and -GRET are 31.39 ± 1.04 and 37.22 ± 0.73 min, respectively; the AUCs are 0.35 ± 0.008 and 0.018 ± 0.003 RLU·min/particles, respectively. (**g**) BRET-FL half-life and AUC of EP-PalmGRET and -GRET are 28.73 ± 0.98 and 34.17 ± 0.93 min, respectively; the AUCs are 0.739 ± 0.017 and 0.038 ± 0.006 RFU·min/particles, respectively. *, p< 0.05; ****, p<0.0001 with two-tailed Student’s t-test with three independent experiments. **(h)** Western blot analysis of 1.2 μm-filtered EPs from HCA1-GRET/-PalmGRET cells. Black square and red square indicate the samples were processed with and without 50 mM DTT reducing agent, respectively. Expected size: GRET (46 kDa); PalmGRET (49 kDa); Alix (96 kD); CD63 (26 kDa); CD81 (26 kDa).

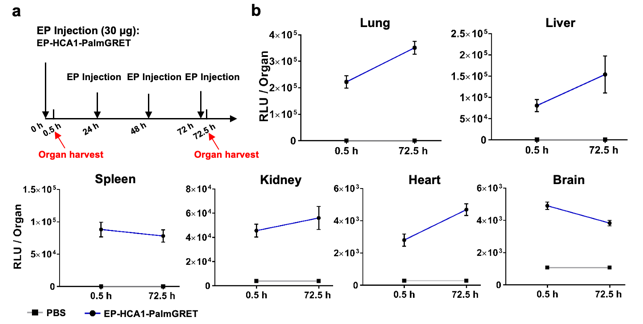

**Figure S9. EP-HCA1-PalmGRET is lung-tropic. (a)** Schematic for EP-tropism experiment. C3H mice were IV-administered with EP-HCA-1-PalmGRET (30 μg) at 0, 24, 48 and 72 h. The organs were harvested at 0.5 and 72.5 h for EP biodistribution analysis. **(b)** Biodistribution analysis indicates a prominent lung-tropism of EP-HCA-1-PalmGRET when compared to other major organs before (0.5 h) and following EP education (72.5 h). PBS was administered in equal volume as a control. *N* = 3 mice per group with technical triplicates.

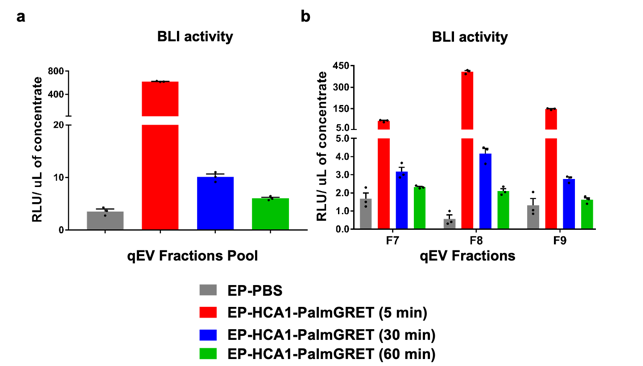

**Figure S10. PalmGRET reports sEV circulation time in C3H immunocompetent mice.** 30 μg EP-HCA-1-PalmGRET was IV-injected into C3H mice’s tail veins. At 5, 30 and 60 min post-EP injection, the blood was collected to isolate the plasma with EDTA as an anti-coagulant. The blood of PBS injected mice was collected at 5 min post-EP injection. Using qEVoriginal 70 nm columns to isolate sEVs from the plasma, fractions Nos. 7–9 were collected for BLI activity assay. BLI activities are shown for **(a)** fraction pools and **(b)** individual fractions. The BLI activities of the fractions were normalized to the volume of concentrate following Amicon Ultra-0.5 with a 10-kDa molecular weight cut-off concentration. *N* = 1 mouse per time point with technical triplicates.

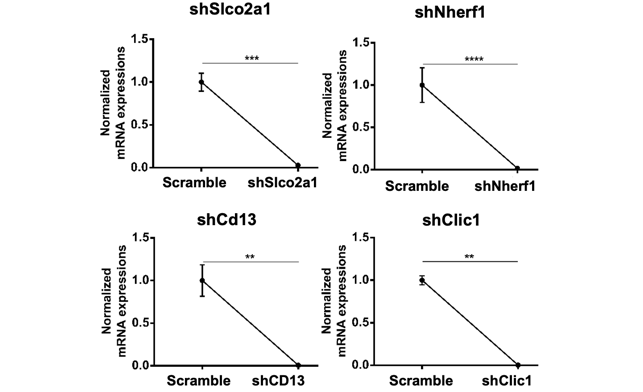

**Figure S11. Real-time quantitative polymerase chain reaction (RT-qPCR) of HCA1-PalmGRET cell with gene silencing of lung-tropic protein candidates.** RT-PCR charts showing >90% gene silencing efficiency of lung-tropism protein candidates (shSlco2a1, shNherf1, shCD13, shClic1) in HCA1-PalmGRET cell. β-actin was used as internal control and the charts were normalized to Scramble. **, p< 0.01; ***, p< 0.001;****, p<0.0001 with two-tailed Student’s t-test with technical triplicate.

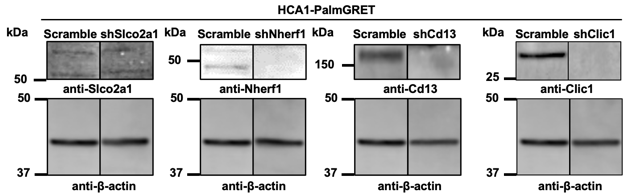

**Figure S12. Western blots demonstrating shRNA-mediated knockdown of lung-tropic protein candidates in HCA1-PalmGRET cell.** Slco2a1 (65 kDa), Nherf1 (40 kDa), Cd13 (110 kDa), Clic1 (27 kDa) expressions of HCA1-PalmGRET cells were significantly reduced by respective shRNAs. β-actin (42 kDa) was as a loading control.

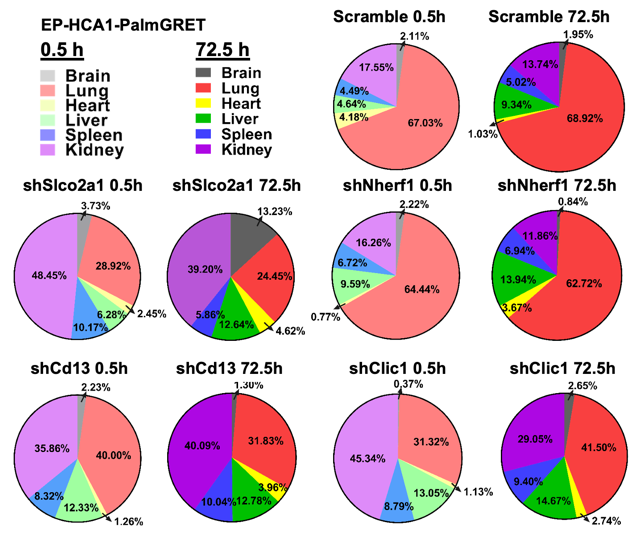

**Figure S13.** **Biodistribution analysis of EP-HCA1-PalmGRET demonstrates dynamic EP distribution following redirected EP lung-tropism.** Pie charts illustrating altered EP biodistributions in immunocompetent C3H mice following Slco2a1, Cd13 and Clic1 knockdowns of EP-HCA1-PalmGRET when compared with the Scramble control. Proportions for each organ at different time points were acquired by$\frac{Mean RLU (Organ)}{\Sigma of Mean ELU(All organs)}$. *N* = 3 mice per group with technical triplicates.

**Table S1. Primer list for cloning**

| Primer name | Sequence (5' → 3') |
| --- | --- |
| GpNluc-Fwd | AAAAAAGCAGGCTCGAGCCACCATGGTGAGCAAGGGC |
| GpNluc-Rev | AGAAAGCTGGGTCTAGAATTACGCCAGAATGCGT |
| PalmGpNluc-Fwd-1 | ATGCTGTGCTGTATGAGAAGAACCAAACAGGTTGAAAAGA  ATGATGAGGACCAAAAGATCATGGTGAGCAAGGGC |
| PalmGpNluc-Rev-1 | TTACGCCAGAATGCGT |
| PalmGpNluc-Fwd-2 | TACAAAAAAGCAGGCTCCACCATGCTGTGCTGTATGAGAA |
| PalmGpNluc-Rev-2 | AGAAAGCTGGGTCTAGAATTACGCCAGAATGCGT |

**Table S2. Antibody list**

| **Name** | **Isotype** | **Host** | **Dilution for WB** | **Dilution for Dot blot** | **Dilution for IHC** | **Vendor** | **Cat. No.** |
| --- | --- | --- | --- | --- | --- | --- | --- |
| GFP | IgG | Rabbit | 1/ 3,000 | 1/ 3,000 | N/A | GeneTex | GTX113617 |
| GFP | IgY | Chicken | N/A | N/A | 1/ 700(lung); 1/ 250(liver); 1/ 700(spleen); 1/ 500 (kidney) | GeneTex | GTX13970 |
| CD63 | IgG | Rabbit | 1/ 500 | N/A | N/A | Abcam | ab216130 |
| CD63 | IgG | Mouse | 1/ 500 | N/A | N/A | Abcam | ab59479 |
| CD81 | IgG | Rabbit | 1/ 1,000 | N/A | N/A | Cell Signalling Technology | 10037S |
| CD81 | IgG | Mouse | 1/ 1,000 | N/A | N/A | Abcam | ab79559 |
| ALIX | IgG | Mouse | 1/ 400 | N/A | N/A | Santa Cruz | sc53540 |
| GAPDH | IgG | Rabbit | 1/ 3,000 | N/A | N/A | Novus | NB300-221 |
| β-actin | IgG | Mouse | 1/ 3,000 | N/A | N/A | Novus | NB600-501-0 |
| CD13(ANPEP) | IgG | Rabbit | 1/ 500 | N/A | N/A | Abcam | ab108310 |
| CLIC1 | IgG | Rabbit | 1/ 500 | N/A | N/A | Proteintech | 14545-1-AP |
| NHERF1 | IgG | Rabbit | 1/ 1,000 | N/A | N/A | Abcam | ab3452 |
| SLCO2A1 | IgG | Rabbit | 1/ 500 | N/A | N/A | Proteintech | 14327-1-AP |
| Rabbit-HRP | IgG | Goat | 1/ 15,000 (sucrose gradient) | 1/ 15,000 | N/A | Jackson ImmunoResearch | 111-035-003 |
| Mouse-HRP | IgG | Goat | 1/ 5,000 (sucrose gradient) | N/A | N/A | Jackson ImmunoResearch | 115-035-003 |
| Rabbit-IRDye 800CW | IgG | Goat | 1/ 5,000 | N/A | N/A | Abcam | ab216773 |
| Mouse-IRDye 680RD | IgG | Goat | 1/ 5,000 | N/A | N/A | LiCoR | 926-68070 |
| Chicken-Alexa Fluor 568 | IgG | Goat | N/A | N/A | 1/ 1,000(lung); 1/ 2,500 (liver); 1/ 1,500 (spleen); 1/ 2,000 (kidney) | Thermofisher | A-11041 |

**Table S3. Examined Cytokine list of mice serum for post-EV administration**

| **Sample Name** | **Murine IL-1β** | | **Murine IL-2** | | **Murine IL-4** | | **Murine IL-5** | | **Murine IL-6** | | **Murine IL-10** | | **Murine IL-12 (P35)** | | **Murine IL-13** | | **Murine IL-17A** | | **Murine IL-22** | | **Murine IL-23(P19)** | | **Murine TNF-α** | | **Murine IFN-γ** | |
| --- | --- | --- | --- | --- | --- | --- | --- | --- | --- | --- | --- | --- | --- | --- | --- | --- | --- | --- | --- | --- | --- | --- | --- | --- | --- | --- |
| **C3H-blank 1** | OOR < | OOR < | *2.60 | *2.60 | *0.47 | *0.43 | OOR < | OOR < | OOR < | OOR < | OOR < | OOR < | *1.52 | *2.66 | OOR < | OOR < | OOR < | OOR < | *0.08 | OOR < | OOR < | OOR < | OOR < | OOR < | *11.50 | *11.94 |
| **C3H-blank 2** | OOR < | OOR < | *3.11 | *1.54 | *0.47 | *0.47 | OOR < | OOR < | OOR < | OOR < | OOR < | OOR < | *1.34 | *1.71 | OOR < | OOR < | OOR < | OOR < | *3.59 | *2.47 | OOR < | OOR < | OOR < | OOR < | *12.39 | *11.50 |
| **C3H-blank 3** | OOR < | OOR < | *9.24 | *9.24 | *0.56 | *0.60 | OOR < | OOR < | OOR < | OOR < | OOR < | OOR < | *2.08 | *2.08 | OOR < | OOR < | OOR < | OOR < | OOR < | OOR < | OOR < | OOR < | OOR < | OOR < | *11.06 | *10.62 |
| **0.5h EV-HCA-1-PalmGp 1** | OOR < | OOR < | OOR < | OOR < | *0.34 | *0.91 | OOR < | OOR < | 18.25 | 17.99 | OOR < | OOR < | *2.08 | *2.46 | OOR < | OOR < | OOR < | OOR < | OOR < | OOR < | OOR < | OOR < | *28.05 | *43.56 | *12.84 | *12.84 |
| **0.5h EV-HCA-1-PalmGp 2** | OOR < | OOR < | OOR < | OOR < | *0.56 | *0.69 | OOR < | OOR < | 18.51 | *16.15 | OOR < | OOR < | *1.89 | *1.89 | OOR < | OOR < | OOR < | OOR < | OOR < | OOR < | OOR < | OOR < | *28.05 | *15.86 | *12.17 | *11.06 |
| **0.5h EV-HCA-1-PalmGp 3** | OOR < | OOR < | OOR < | OOR < | *0.65 | *0.82 | OOR < | OOR < | 68.14 | 55.92 | OOR < | OOR < | *1.71 | *1.34 | OOR < | OOR < | *5.39 | OOR < | OOR < | OOR < | OOR < | OOR < | 60.8 | *49.92 | *11.94 | *10.62 |
| **24.5h EV-HCA-1-PalmGp 1** | OOR < | OOR < | OOR < | OOR < | *0.47 | *0.20 | OOR < | OOR < | OOR < | OOR < | OOR < | OOR < | *2.08 | *1.71 | OOR < | OOR < | OOR < | OOR < | OOR < | OOR < | OOR < | OOR < | *33.18 | *11.92 | *10.62 | *9.75 |
| **24.5h EV-HCA-1-PalmGp 2** | OOR < | OOR < | OOR < | *4.59 | *0.65 | *0.56 | OOR < | OOR < | OOR < | OOR < | OOR < | OOR < | *2.08 | *2.46 | OOR < | OOR < | OOR < | OOR < | OOR < | OOR < | OOR < | OOR < | OOR < | *35.60 | *14.64 | *14.19 |
| **24.5h EV-HCA-1-PalmGp 3** | OOR < | OOR < | *0.98 | *2.60 | *0.65 | *0.51 | OOR < | OOR < | *5.95 | *4.85 | OOR < | OOR < | *2.46 | *2.46 | OOR < | OOR < | OOR < | OOR < | OOR < | OOR < | OOR < | OOR < | *22.41 | *22.41 | *17.41 | *15.56 |
| **48.5h EV-HCA-1-PalmGp 1** | OOR < | OOR < | OOR < | OOR < | *0.82 | *0.29 | OOR < | OOR < | *4.66 | *3.45 | OOR < | OOR < | *2.08 | *2.08 | OOR < | OOR < | OOR < | OOR < | OOR < | OOR < | OOR < | OOR < | *30.67 | *6.87 | *13.74 | *11.94 |
| **48.5h EV-HCA-1-PalmGp 2** | OOR < | OOR < | OOR < | OOR < | *0.51 | *0.38 | OOR < | OOR < | OOR < | OOR < | OOR < | OOR < | *1.52 | *1.71 | OOR < | OOR < | OOR < | OOR < | OOR < | OOR < | OOR < | OOR < | *30.67 | *26.70 | *16.48 | *13.74 |
| **48.5h EV-HCA-1-PalmGp 3** | OOR < | OOR < | OOR < | OOR < | *0.51 | *0.29 | OOR < | OOR < | *11.16 | *9.22 | OOR < | OOR < | *2.08 | *1.16 | OOR < | OOR < | OOR < | OOR < | OOR < | OOR < | OOR < | OOR < | *44.65 | *30.67 | *14.19 | *13.74 |
| **72.5h EV-HCA-1-PalmGp 1** | OOR < | OOR < | OOR < | OOR < | *0.47 | *0.29 | OOR < | OOR < | OOR < | OOR < | OOR < | OOR < | *1.71 | *1.71 | OOR < | OOR < | OOR < | OOR < | OOR < | OOR < | OOR < | OOR < | OOR < | OOR < | *11.72 | *13.74 |
| **72.5h EV-HCA-1-PalmGp 2** | OOR < | OOR < | OOR < | OOR < | *0.29 | *0.38 | OOR < | OOR < | OOR < | OOR < | OOR < | OOR < | *1.71 | *0.63 | OOR < | OOR < | OOR < | OOR < | OOR < | OOR < | OOR < | OOR < | *6.87 | OOR < | *11.94 | *11.94 |
| **72.5h EV-HCA-1-PalmGp 3** | OOR < | OOR < | OOR < | OOR < | *0.56 | *0.47 | OOR < | OOR < | OOR < | OOR < | OOR < | OOR < | *1.34 | *2.08 | OOR < | OOR < | OOR < | OOR < | OOR < | OOR < | OOR < | OOR < | OOR < | OOR < | *12.84 | *12.39 |

Each sample was test in duplicates;

*Value = Value extrapolated beyond standard range;

OOR< = Out of Range Below;

Concentration Units = pg/ml

**Table S4. Analyzed proteomic data of EV-HCA-1-PalmGp**

**Please attached file.**

**Table S5. Target sequence of shRNAs for knockdown experiment**

| Name | Target sequence |
| --- | --- |
| Slco2a1 | GCCTATGCCAACTTACTCATT |
| Cd13 | CCTTTCTGTTATCCCTGTCAT |
| Clic1 | GCCCTGAAGGTTCTAGACAAT |
| Nherf1 | CGATACCAGTGAGGAGCTAAA |
| Scramble | N/A |

**Table S6. Primers for RT-qPCR of knockdown experiment**

| Name | Forward | Reverse |
| --- | --- | --- |
| Slco2a1 | TCTCCACGTTCCTCAACAAGT | GCAGAGGGAAAACAAAACGCT |
| Cd13 | ACCAGAGTGCAAAGTTCCAGA | CCAGGTTGAAGGAGTCGTGG |
| Clic1 | AATCAAACCCAGCACTCAATG | CAGCACTGGTTTCATCCACTT |
| Nherf1 | CCTCCAGCGATACCAGTGAG | CACAGCCAAGGAGATGTTGAG |
| β-Actin | TCCATCATGAAGTGTGACGT | TACTCCTGCTTGCTGATCCACAT |

**Movie S1. 3D reconstruction of live-cell confocal Z-stack images of 293T-PalmGRET and 293T-PalmtdTomato co-culture.**

**Movie S2. Live-cell SRRF nanoscopy of 293T-PalmGRET cells.**

**Movie S3. Boxed region of Movie S2 showing budding-like protrusion.**
